# An insect brain-based bioelectronic neural sensor for the systemic detection and precise classification of endometriosis

**DOI:** 10.1101/2025.06.03.657650

**Authors:** Simon W. Sanchez, Erin L. Vegter, Michael Parnas, Yong Song, Asgerally T. Fazleabas, Debajit Saha

## Abstract

Endometriosis is a chronic inflammatory disease with limited screening options and a recognized diagnostic delay. To investigate detection potentialities, a bioelectronic sensor is realized to detect endometriotic vs endometrial models via their emitted volatile organic compounds (VOCs) by leveraging an insect olfactory system combined with computational analytical techniques for classification. Our analyses of cell culture headspace-evoked neural responses show that our sensor can distinguish multiple cell lines by their ‘scent’ (i.e., emitted VOC mixture). By combining neural responses across experiments, we obtained high-dimensional population neural response templates that were used to classify unknown samples with a high accuracy. We obtained an accuracy of 89% in differentiating 4 cell lines at two growth timepoints (24 hr. and 72 hr.) and obtained an accuracy of 88% in classifying epithelial co-cultures of endometriotic and endometrial cell lines cultured at 0%, 25%, 50%, 75%, and 100% in ascending/descending ratios. Our results support the hypothesis that endometriosis is detectable via the metabolic differences found in the emitted gas mixtures or ‘scent’ from various cell lines and demonstrates the effectiveness of our sensor in distinguishing the subtle changes in endometriotic vs endometrial models pertaining to durations in growth and co-cultures.

## Introduction

Endometriosis is a chronic gynecological disease characterized by the presence of ectopic endometrial-like tissue outside the uterus ^1^. This inflammatory disease affects approximately 10% of women of reproductive age and is clinically associated with a range of symptoms that are common to other gynecological and non-gynecological disorders ^2–7^. Women with endometriosis experience a substantial decrease in quality of life and the financial and emotional burden of endometriosis is amplified by the diagnostic delay which ranges from 4 to 11 years ^8–13^. Barriers to the early diagnosis and treatment of endometriosis are a result of the current standard of surgical diagnosis via a diagnostic laparoscopy with histological verification to confirm the presence of endometriotic glands and stroma ^14,15^. Thus, a shift from surgical diagnosis to a clinical one is imperative and should be explored ^16–18^.

Extensive efforts have been made to find specific biomarkers for endometriosis however these biomarkers have not reached clinical validation due to confounding factors hindering progress such as inadequate evidence and variability across research studies ^19–26^. Due to the systemic nature of the disease, changes in the cardiovascular, neurological, immune, and metabolic function have been well established and leads to the question of whether a multiplexed holistic approach utilizing the systemic processes of the disease can be used for diagnosis ^5,6,27^. Here, we propose a novel concept that pathogenic and control endometrial cells generate distinct profiles of volatile organic compounds (VOCs) due to the changes in metabolic processes which can be employed for detection. VOCs are a diverse group of carbon-based chemicals that vaporize readily due to their relatively high vapor pressure at room temperature and atmospheric pressure^28^. VOCs are emitted from the body via exhaled breath and are also found in biological fluids such as blood, urine, feces, and sweat signaling the systemic processes contributing to these compounds ^29,30^. Distinct patterns of VOCs are based on the concept that pathological processes, happening as a result of the disease, can alter the relative concentrations of existing VOCs ^31–34^. The presence of several diseases including cancer changes the metabolism and the metabolite composition in the cell, in its’ surrounding medium, and as a result in the emitted VOC profile found in the headspace of *in-vitro* cultured cells ^35–42^. Additionally, these VOC profiles have been found in exhaled human breath using chemical analysis (e.g., gas chromatography-mass spectrometry) and differences in the VOC composition have been found between control and cancer patients ^43–49^. For a systemic and complex disease that is endometriosis, a chemical gas sensor for generalizable and nonspecific odor recognition would be essential to achieve such detection capabilities.

Biological systems have solved the problem of chemical sensing via evolution and have converged to a solution that is architecturally and operationally similar across species ^50–52^. This signifies that there might be a superior solution for gas sensing that is still inaccessible from an engineering standpoint ^53^. Living organisms (e.g., canines) use biological olfaction to detect VOC mixtures via their robust odor recognition capabilities and generalization for chemicals across varying concentrations which can be employed for disease detection ^54–59^.

Circumventing the need to detect a single or known panel of biomarkers, we developed an insect-based neural sensor and computational analytical techniques to classify endometriotic vs endometrial cell lines through the use of VOC gas mixtures. The sensor was developed by incorporating electrophysiological recordings of antennal lobe (AL) projection neurons with cross validation techniques across high-dimensional population neural responses. Responses from both single and population neurons differentiated multiple pathogenic and control cell lines including epithelial cells and stromal cells at multiple timepoints. By employing spatiotemporal population responses analyses, multiple timepoints of each cell line could be classified with a high accuracy. Additionally, co-cultures of combined pathogenic and control cells prepared in multiple ratios could be classified indicating the effectiveness of the sensor to detect subtle differences the progression of pathogenic and control mixtures. Overall, our results highlight the detection capabilities of biological olfactory systems, neural data analysis tools, and together put forth a platform for the potential diagnosis of endometriosis using volatile gas mixtures for a systemic and holistic approach.

## Results

### Endometriotic and endometrial cell lines elicit distinct neural responses

We initially sought to investigate the locust AL neural response patterns to endometriotic and endometrial emitted cell culture gas mixtures. Recording the neural responses from the locust AL allows the recording of the chemosensory information as it funnels into the antennal lobe from the locust antenna before diverging onto higher-order structures ^56,60–66^. Using the experimental setup schematically shown in Fig. 1a, we delivered the volatile chemical gas mixtures contained within a cell culture flask to the locust antenna via an olfactometer system and neural responses were recorded from the locust AL. Using this system, we are able to control the timing and volume of gas mixture delivery to the locust antenna while concurrently performing *in vivo* extracellular recordings (see Methods). Two endometriotic cell lines (12Z; epithelial and iEc-ESC; stromal) and two endometrial cell lines (H1657-iEEC; epithelial and H1644-iESC; stromal) were grown in identical cell culture media after seeding each cell line at the same initial cell number (Fig. 1b) ^67,68^. All four cell lines were grown in individual sterile airtight T-25 flasks for 72 hours before *in vivo* extracellular neural recordings. Volatile chemical gas mixtures emitted from the cell culture and a media control were then delivered in precise amounts to the locust antenna for 4 seconds (Fig. 1a). Simultaneously, *in vivo* extracellular neural recordings were obtained from the locust antennal lobe.

**Fig. 1:**
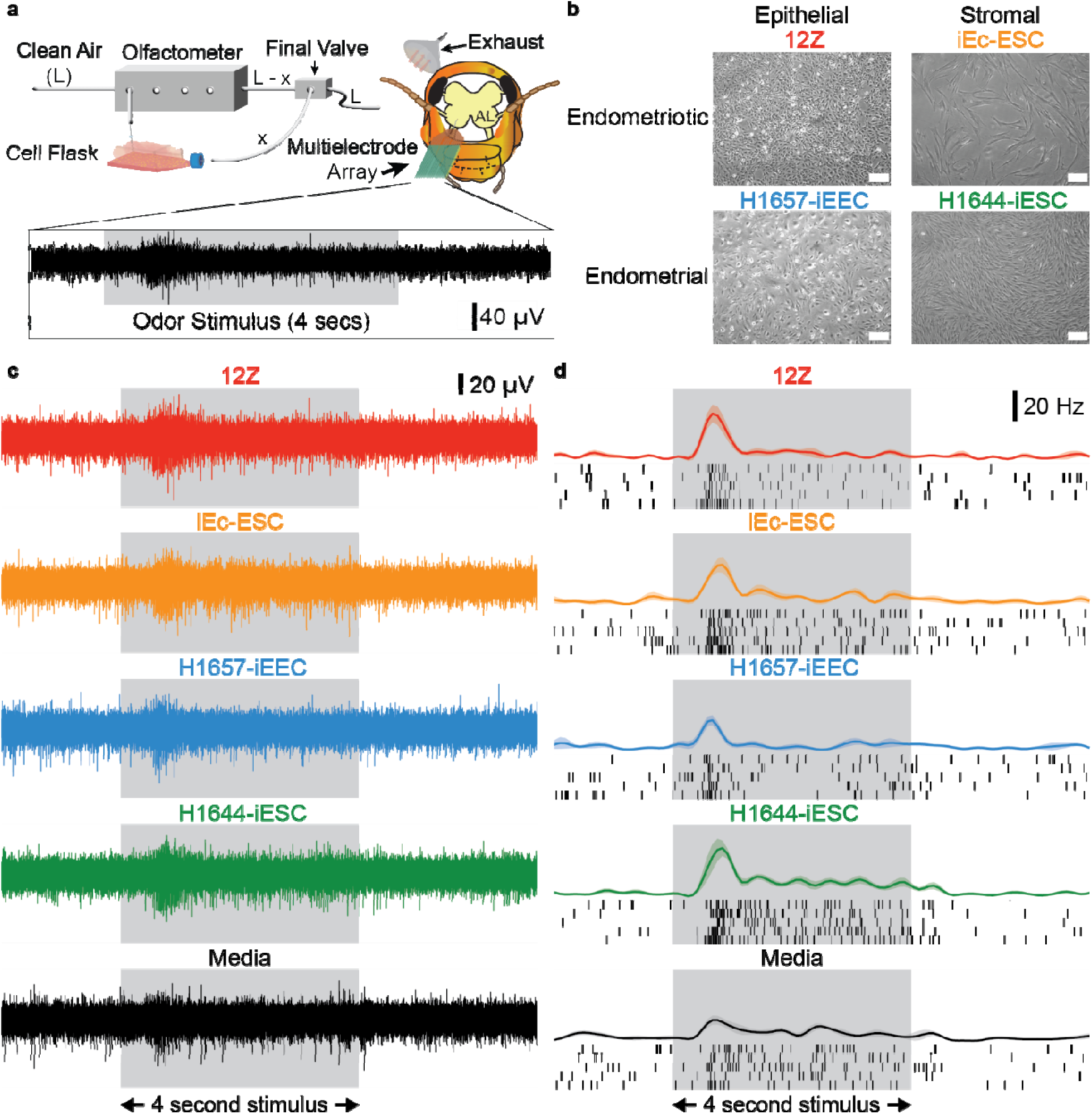
Distinct olfactory neural responses to endometriotic and endometrial cell lines. **a** Schematic of experimental setup. In this setup, cell culture headspace gas mixtures are delivered to the chemosensory array (antenna) and neural responses are recorded from the antennal lobe region of the locust brain using a multielectrode array. A multielectrode array placed within the locust antennal lobe records the cell culture headspace evoked neural responses. At the bottom, a representative peri-stimulus voltage trace (PSVT) is shown with a 4 second odor presentation window in gray. **b** Representative images of the four cell cultures used in this study. Immortalized human ectopic endometriotic epithelial cells (12Z) and stromal cells (iEc-ESC) are shown on the top. Immortalized endometrial epithelial cells (H1657-iEEC) and stromal cells (H1644-iESC) are shown on the bottom. All the images are shown at 72-hour post-seeding. The white scale bar indicates 100 µm. **c** Representative extracellular cell culture headspace-evoked neural voltage responses from a single location within the locust antennal lobe is shown. The grey box indicates the 4 second stimulus presentation window. **d** Distinct neural spiking responses to all cell culture and media headspace gas mixtures is shown. Representative raster plots from one neuron spike sorted from the voltage trace shown in panel c shows spiking patterns from five trials of each cell line and media. Each black line in the raster plot indicates an action potential, or spiking event, from the neuron. Peri-stimulus time histograms (PSTHs) are shown above each raster plot for all cell cultures and media. Trial-averaged PSTHs are plotted with the shaded region indicating the standard error of the mean (SEM). The gray box indicates the 4 second stimulus presentation window.

From these recordings, we observed that the volatile chemical gas mixtures emitted from each of the cell cultures and media elicited changes in the neural spiking responses in the locust AL. The increase in voltage or ‘thickening’ of the voltage trace soon after the volatile gas mixture was delivered to the locust antenna indicated a stimulus-evoked response (Fig. 1c). To validate the stimulus-evoked response, the single unit activity of a neuron was obtained via spike sorting (see Methods) following previously published methods from the same voltage trace shown in Fig. 1c ^69^. For the single representative neuron, the firing rate was plotted using raster plots and peri-stimulus time histograms (PSTHs) displaying distinct neural responses to each cell line emitted gas mixture and media (Fig. 1d). Similar to the voltage trace an increase in spiking activity can be seen soon after the volatile gas mixture was delivered to the locust antenna. Here, we can begin to notice the subtle changes in neural spiking frequency to each cell line’s emitted gas mixture over the 4 second stimulus presentation. In total we conducted 36 electrophysiological recordings, and from these recordings we spike sorted a total of 58 neurons. To validate the differences between different stimulus conditions (i.e., gas mixtures from different cell lines), we performed a one-way ANOVA with Bonferroni correction analysis comparing the spike counts to all combinations of two stimulus conditions from our cell line panel (p < 0.05, d.f. = 4, 20, one-way ANOVA with Bonferroni correction; Supplementary Fig. 1). To do this we compared the total spike counts over the 4 second stimulus averaged for all 5 trials for every combination of two stimulus conditions. From this, we see that for every combination, we have several neurons that show significant difference with increased (plotted in red) or decreased (plotted in blue) spike counts relative to the stimulus condition on the X-axis (Supplementary Fig. 1). A neuron that did respond significantly different was plotted in gray. The ability of individual neurons to have varied and significantly different responses to various endometriotic and endometrial cell lines showcases the distinction of gas mixtures emitted from these cell lines at the individual neuron level (Fig. 1 and Supplementary Fig. 1). While single neuron level differentiation is impressive for multiple cell lines, odor identity is thought to be encoded at the population level and further analysis is required to assess this.

### Spatiotemporal neural responses classify endometriotic and endometrial cell lines

In the antennal lobe, it is thought that the identity of an odor is not encoded by a single PN, but by a population of PNs via a time-varying spatiotemporal response ^60,70–74^. Consequently, we next examined the neural population response to each emitted gas mixture from all cell lines and media over a specified time window capturing the evolution of the population response over time (i.e., spatiotemporal response). Before proceeding solely with the population of spike sorted neurons (n = 58) for analysis, voltage traces obtained from all extracellular neural recordings (n = 36) were processed through a root mean squared (RMS) filter that allowed us proceed with two analyses in parallel (Supplementary Fig. 2). Using an RMS filter for data processing is a computationally inexpensive technique that is automated and fast in comparison to manual spike sorting that requires supervision from skilled personnel and is known to have user performance variability (Supplementary Table 1) ^75,76^. An RMS-based data processing technique has the potential be clinically useful for rapid processing and classification of samples for a disease (i.e., endometriosis) with a known diagnostic latency ^77^. However, it should be noted that RMS filtering does not allow for single unit activity to be obtained as with spike sorting that eliminates signals that do not pass statistical tests ^69^. A representative voltage trace that has been filtered with the RMS transformation is shown in Supplementary Fig. 3.

Proceeding forward with the RMS filtered data, we used dimensionality reduction techniques (i.e., principal component analysis; PCA and linear discriminant analysis; LDA) to reduce the data from a high dimensional dataset (i.e., 36 dimensions) to a low dimensional subspace (see Methods). This allowed us to visualize the population neural responses in 3-dimensional PCA and LDA subspace as shown in Fig. 2a and 2b, respectively. Within 3-dimensional PCA subspace, we can qualitatively identify distinct and independent trajectories that represent the population neural responses to the emitted gas mixture of each of the cell lines and media over 1.25 seconds during the stimulus presentation window (Fig. 2a). The unique paths and separate angles in which these trajectories move and diverge signify that these cell lines and media are eliciting distinct responses at the population level (Fig. 2a). Additionally, using LDA, we can again qualitatively identify distinct and separate clusters of 50 millisecond time bins representing the 1.25 seconds (25 total time bins) during the stimulus presentation window for each cell line and media (Fig. 2b). Furthermore, using our parallel analysis with the population of spike sorted neurons, unique trajectories and separate clusters are also obtained in PCA and LDA subspace (Supplementary Fig. 4a and b). Qualitatively these results confirm, at the population level, the ability of a population of antennal lobe PNs to differentiate endometriotic and endometrial cell lines including media by their emitted gas mixture, however a quantitative analysis is still needed to measure the classification accuracy of this system.

**Fig. 2:**
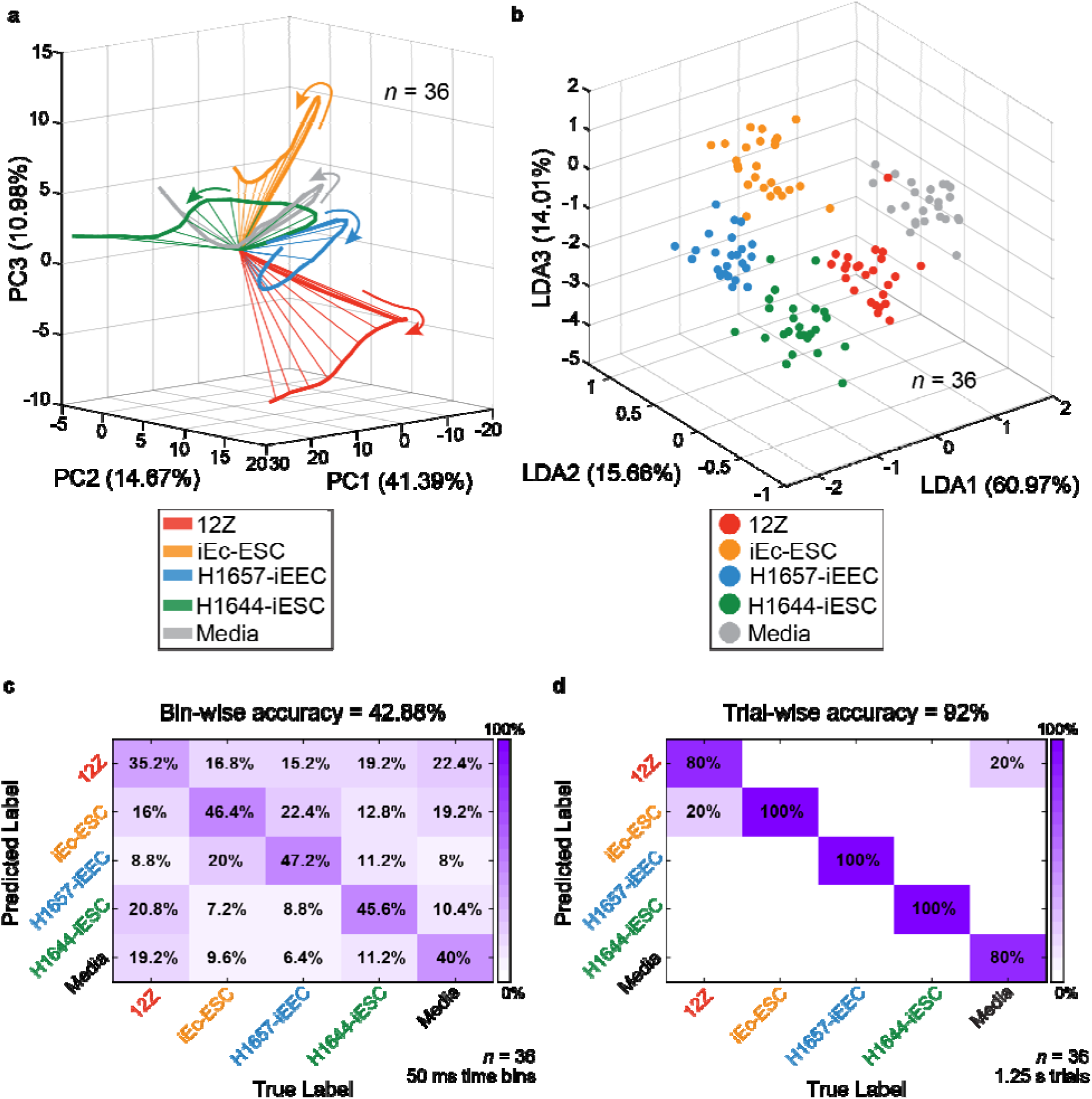
Spatiotemporal population neural responses classify endometriotic and endometrial cell lines. **a** Dimensionality reduction of high dimensional neural response vectors via principal component analysis (PCA) shows distinct temporal neural trajectories in the PCA subspace. Arrows specify the direction of the neural trajectories from the origin which indicates the start of the time window. The time window plotted is 0.5 – 1.75 seconds after stimulus onset. **b** Dimensionality reduction of high dimensional neural response vectors via linear discriminant analysis (LDA) shows distinct neural response clusters in the LDA subspace. Each point in the LDA subspace indicates a 50-millisecond time bin within the time window of 0.5 – 1.75 seconds after stimulus onset. **c** Confusion matrix summarizing the high dimensional leave-one-trial-out (LOTO) analysis for the classification of 50 millisecond time bins within the time window of 0.5 – 1.75 seconds after stimulus onset. **d** Confusion matrix summarizing the high dimensional LOTO analysis for the classification of 1.25 second trials for the entire time window of 0.5 – 1.75 seconds after stimulus onset. Data was RMS filtered (see Methods) from 36 electrophysiological recording locations for analysis.

In order to achieve a quantitative measure for the classification accuracy of this system, we next sought to predict the classification success of unknown neural templates for each of the cell lines and media using a high-dimensional matching analysis with neural training templates denoted leave-one-trial-out analysis (LOTO) (see Methods). A schematic representation of this analysis has been previously described ^57^. Neural responses were divided into 50 millisecond time bins due to the 20 Hz local field oscillation observed in the locust antennal lobe for analysis ^78–80^. The results from this bin-wise LOTO analysis are summarized in a confusion matrix shown in Fig. 2c where we see that most of the time bins are being classified correctly indicated by the dark diagonal values going down the matrix with a total accuracy of 42.88%. To extend on this analysis, we took a trial-wise approach by assigning a whole trial in a winner takes all scenario to a sample determined by the mode of the time bins for each trial (Fig. 2d). With this trial-wise approach, we obtained a 92% classification accuracy (Fig. 2d). The spike sorted classification results display similar classification accuracies of 41.44% and 94% using bin-wise and trial-wise methods, respectively (Supplementary Fig. 4c and d). By taking advantage of the spatiotemporal population PN responses in the locust antennal lobe, differentiation and classification of emitted gas mixtures from endometriotic and endometrial cell lines is attained.

### Cell growth duration influences neural response patterns

To further demonstrate the reliability and repeatability of our sensor, we next investigated whether individual neurons would elicit distinct responses to the same cell lines used previously but at multiple timepoints. Representative images of all cell lines are shown in Fig. 3a. Volatile chemical gas mixtures emitted from each of the cell cultures at the two timepoints and media elicited changes in the neural spiking responses in the locust antennal lobe as shown in Fig. 3b. Interestingly, representative neuron 1 showed a consistent response across the endometrial cell lines regardless of the timepoint (24 or 72 hr.) in comparison to the endometriotic cell lines suggesting that this neuron is “tuned” towards the volatile chemicals emitted from the endometrial cells. In comparison, representative neuron 2 showed a consistent response across all the cell lines at the 72-hr. timepoint regardless if the cell line were endometriotic or endometrial. This suggests that this neuron is “tuned” towards the concentration of volatiles emitted from the 72-hour timepoint regardless if the cell line was endometrial or endometriotic. Having two neurons with different suggested tuning characteristics implies a complex odor-evoked response and encoding of odor identity inherent for biological olfaction. Moreover, both of these neurons showcase the importance of recording from a sample of PNs in the locust AL in order to capture a breadth of neural responses to various gas mixtures. To validate the differences between different stimulus conditions, we performed a one-way ANOVA with Bonferroni correction analysis comparing the spike counts of all eight cell lines to media (p < 0.05, d.f. = 8, 36, one-way ANOVA with Bonferroni correction; Supplementary Fig. 5). Here, we see neurons that responded significantly different to the cell lines in comparison to the responses evoked by media (Supplementary Fig. 5). Overall, these results showcase the ability of individual neurons to have diverse and varied responses to cell lines and their emitted gas mixtures at multiple timepoints.

**Fig. 3:**
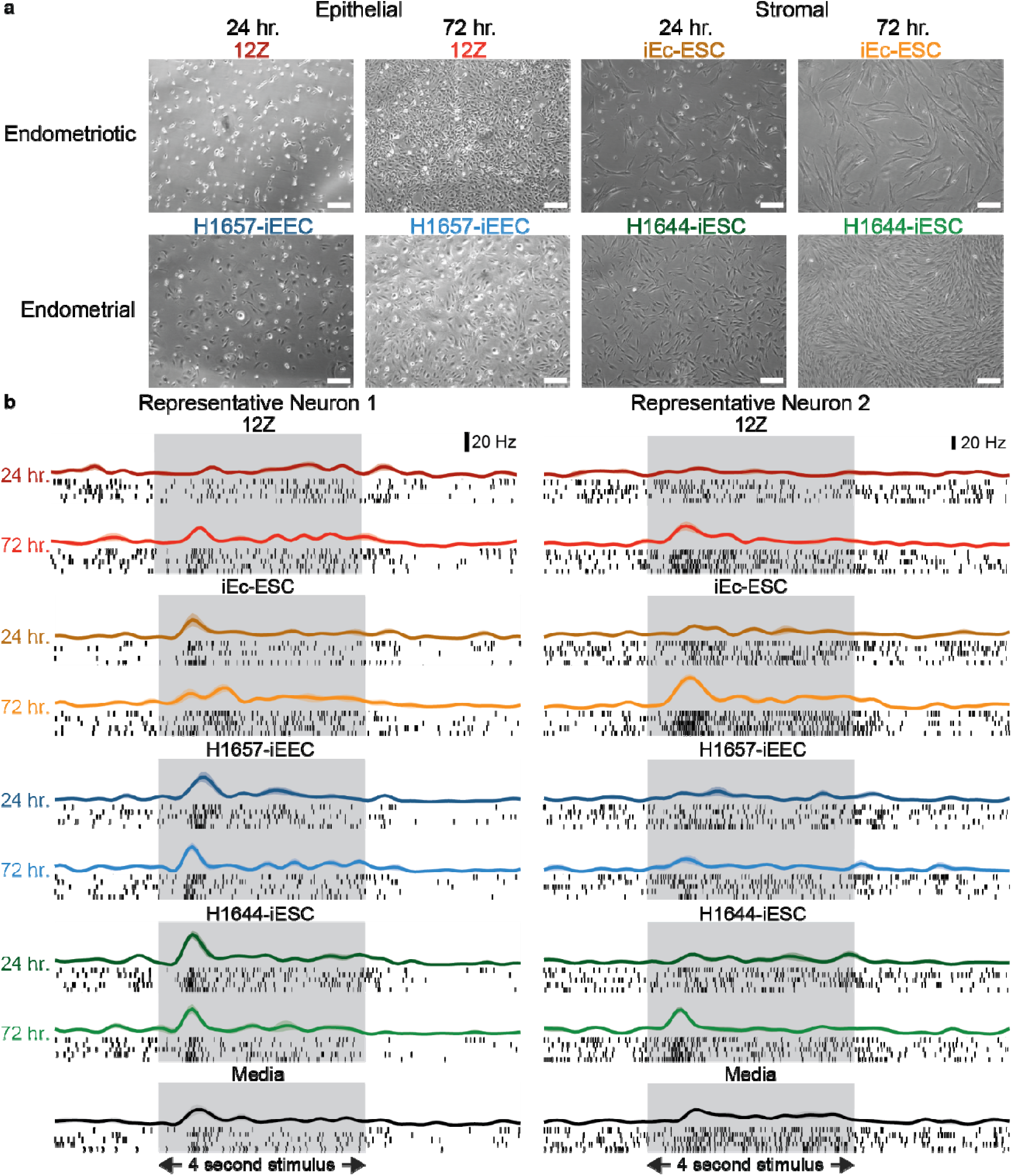
Distinct and tuned neural responses to endometriotic and endometrial cell lines at multiple timepoints. **a** Representative image of the four cell lines at two different timepoints. All the images are shown at either 24 or 72-hour post-seeding. The white scale bar indicates 100 µm. **b** Distinct and tuned neural spiking responses to all cell culture and media headspace gas mixtures is shown for two representative neurons at multiple cell culture time points. Representative raster plots show spiking patterns from five trials of each cell line and media. Each black line in the raster plot indicates an action potential, or spiking event, from the respective neuron. PSTHs are shown above each raster plot. Trial-averaged PSTHs are plotted with the shaded region indicating the SEM. The gray box indicates the 4 second stimulus presentation window.

### Classification of endometriotic and endometrial cell lines at multiple timepoints

To further investigate how the population of neural responses differentiate each of the cell lines and timepoints, we next used dimensionality reduction techniques to qualitatively analyze and visualize our data. To do this we utilized PCA and LDA which allowed us to reduce the dimensionality of our data for visualization purposes (Fig. 4a and b). In PCA subspace we have several trajectories projecting in unique directions with some interesting similarities between the two iEc-ESC cell cultures (24 and 72 hr.) following similar but not exact paths and also interesting distinctions such as the 72 hr. 12Z cell culture extended and separated from all other cell lines (Fig. 4a). In LDA subspace, we see each of the cell line clusters separate although all clusters are situated near each other (Fig. 4b). These dimensionality reduction techniques suggest adequate differentiation within the locust olfactory system for classification. An additional hierarchal clustering analysis was done using the population of spike sorted neurons (Fig. 4c). To do this, we compared the time-matched 50 ms time bins across all trials and cell lines in the experimental panel in high-dimensional space (see Methods). Here, we see two main clusters separated by 72 hr. and 24 hr. cell cultures (Fig. 4c). Furthermore, within these two clusters we also see the separation of endometriotic and endometrial cell lines with only two misclassifications for one trial from the 72 hr. 12Z and 24 hr. H1657-iEEC cell cultures (Fig. 4c). These results suggest that our sensor can distinguish each of the cell lines but also the subtle differences between the cell lines at multiple timepoints.

**Fig. 4:**
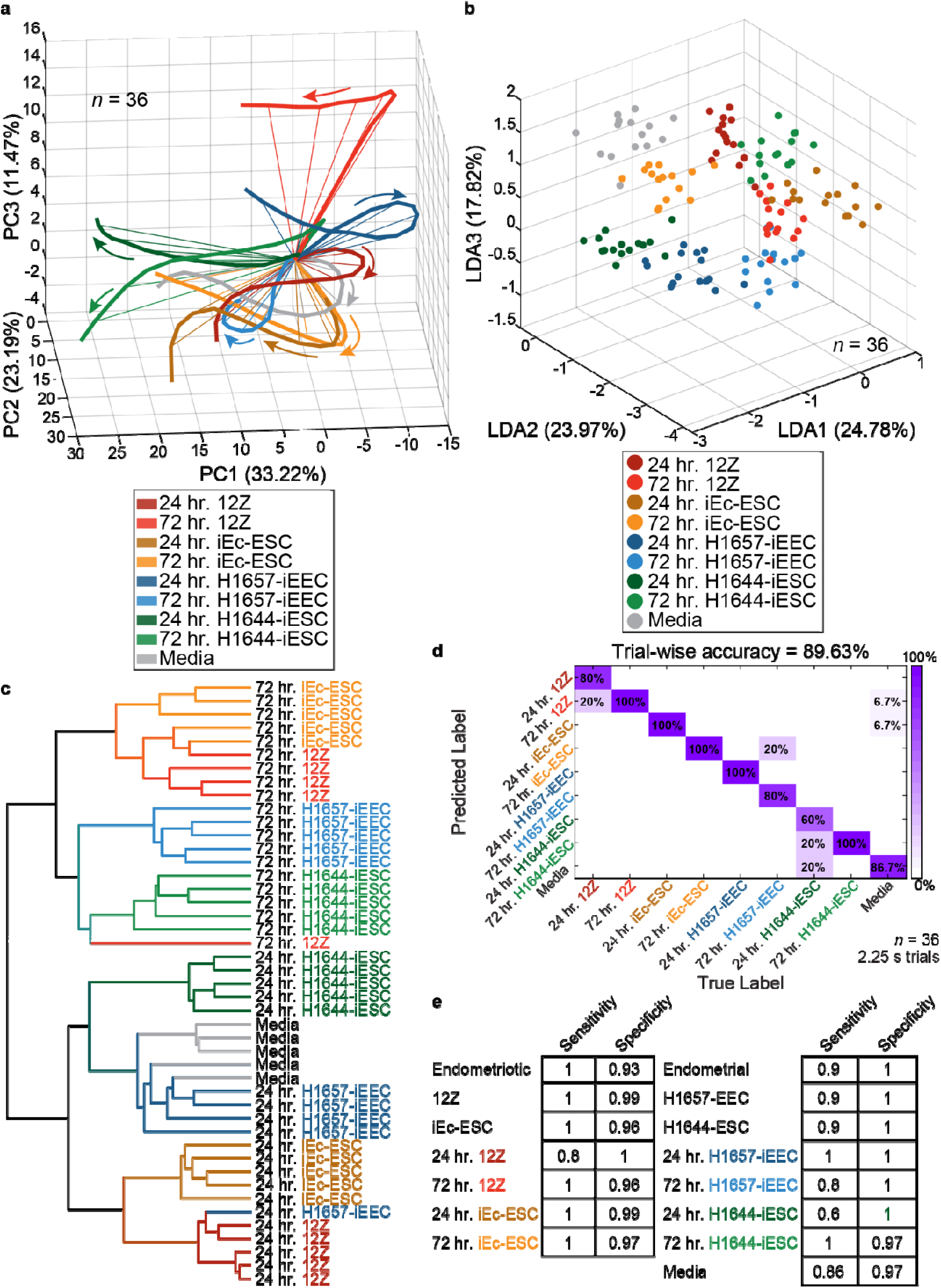
Spatiotemporal population neural responses classify endometriotic and endometrial cell lines at multiple timepoints. **a** Dimensionality reduction of high dimensional neural response vectors via PCA shows distinct temporal neural trajectories in the PCA subspace. Arrows specify the direction of the neural trajectories from the origin which indicates the start of the time window. The time window plotted is 0.5 – 1.25 seconds after stimulus onset. **b** Dimensionality reduction of high dimensional neural response vectors via LDA shows distinct neural response clusters in the LDA subspace. Each point in the LDA subspace indicates a 50-millisecond time bin within the time window of 0.75 – 1.5 seconds after stimulus onset. **c** A hierarchical clustering analysis dendrogram grouped the responses into two clusters separated by the time point (72 hr. and 24 hr.) in which the cells were grown. Moreover, these clusters were further grouped into endometriotic and endometrial clusters within these time points with the exception of a single trial from the cell lines 72 hr. 12Z and 24 hr. H1657-iEEC. The time window for this analysis was 0.5 – 2.25 seconds after the stimulus onset. **d** Confusion matrix summarizing the high dimensional LOTO analysis for the classification of 2.25 second trials for the ‘on’ response time window of 0.25 – 1.5 seconds after stimulus onset in combination with the ‘off’ response time window of 3.5 – 4.5 seconds after stimulus onset (see Supplementary Fig. 6). **e** Sensitivity and specificity table summarizing the results from the confusion matrix in panel d. Data was RMS filtered from 36 electrophysiological recording locations for analysis in panel a, b, and d. Data was spike sorted from 36 electrophysiological recording locations to obtain 58 neurons for analysis in panel c.

To assess the classification accuracy of our sensor we performed a high-dimensional LOTO analysis. Using this analysis, we obtained an accuracy of 89.63% for classifying all cell lines at both timepoints with some notable misclassifications such as the 24 hr. 12Z cell culture classifying as the 72 hr. 12Z cell culture and the 24 hr. H1644-iESC cell culture misclassifying as the 72 hr. H1644-iESC cell culture or media (Fig. 4d). To obtain this high classification accuracy, we combined “on” and “off” response that is displayed during the stimulus presentation (i.e., “on” response) and after the stimulus presentation (i.e., “off” response) (Supplementary Fig. 6). This allowed us to utilize the termination of sensory input for cell line classification and showcases the variability of neural responses to emitted cell culture gas mixtures for these two stimulus modalities ^81,82^. To validate the classification accuracy obtained with our sensor, we applied the LOTO analysis to the 2 seconds before stimulus onset in which baseline neural activity was recorded and obtained an accuracy of 2.59% (Supplementary Fig. 7). Applying the same analyses of dimensionality reduction and LOTO to the population of spike sorted neurons, we obtain similar results (Supplementary Fig. 8a to c). Additionally, the sensitivity and specificity of the sensor was calculated summarizing the results obtained from the LOTO analysis (Fig. 4e and Supplementary Fig. 8d).

### Varying co-culture compositions of endometriotic and endometrial cells are distinguishable

We next examined the ability of our sensor to detect the emitted gas mixtures from both endometriotic (12Z) and endometrial (H1657-iEEC) epithelial cell lines co-cultured within the same flask at different ratios. Representative images are shown in Fig. 5a with the 12Z cell line fluorescently labeled ^83^. In total we used a panel of five cell lines with two of the cultured cell flasks being 100% 12Z and 100% H1657-iEEC and three co-cultured flasks containing varying amounts of each cell line. A flask containing only media was used as a control. A representative neuron is displayed in Fig. 5b showing stimulus-evoked neural responses to all co-cultured cell lines and media. Comparable to the tuned responses we observed for the cell cultures at multiple timepoints, we also observe tuned neural responses to the 100% cell cultures and media as compared to the co-cultured flasks (Supplementary Fig. 9a). Using dimensionality reduction techniques (i.e., PCA and LDA) we observe unique neural trajectories in PCA subspace and clusters in LDA subspace, however in LDA subspace there appears to not be clear separation of two of the co-cultured flasks (50% 12Z 50% H1657-iEEC and 75% 12Z 25% H1657-iEEC) (Supplementary Fig. 9b). Using dimensionality reduction techniques on the population of spike sorted neurons we see similar results (Supplementary Fig. 10a and b). While the high-dimensional hierarchal clustering analysis interestingly shows two main clusters separated by the co-cultures and the individual cell lines (including media) (Supplementary Fig. 10c). These results show that the locust olfactory system possesses a distinctive encoding mechanism for varying ratios of endometriotic and endometrial co-cultures and their emitted gas mixtures.

**Fig. 5:**
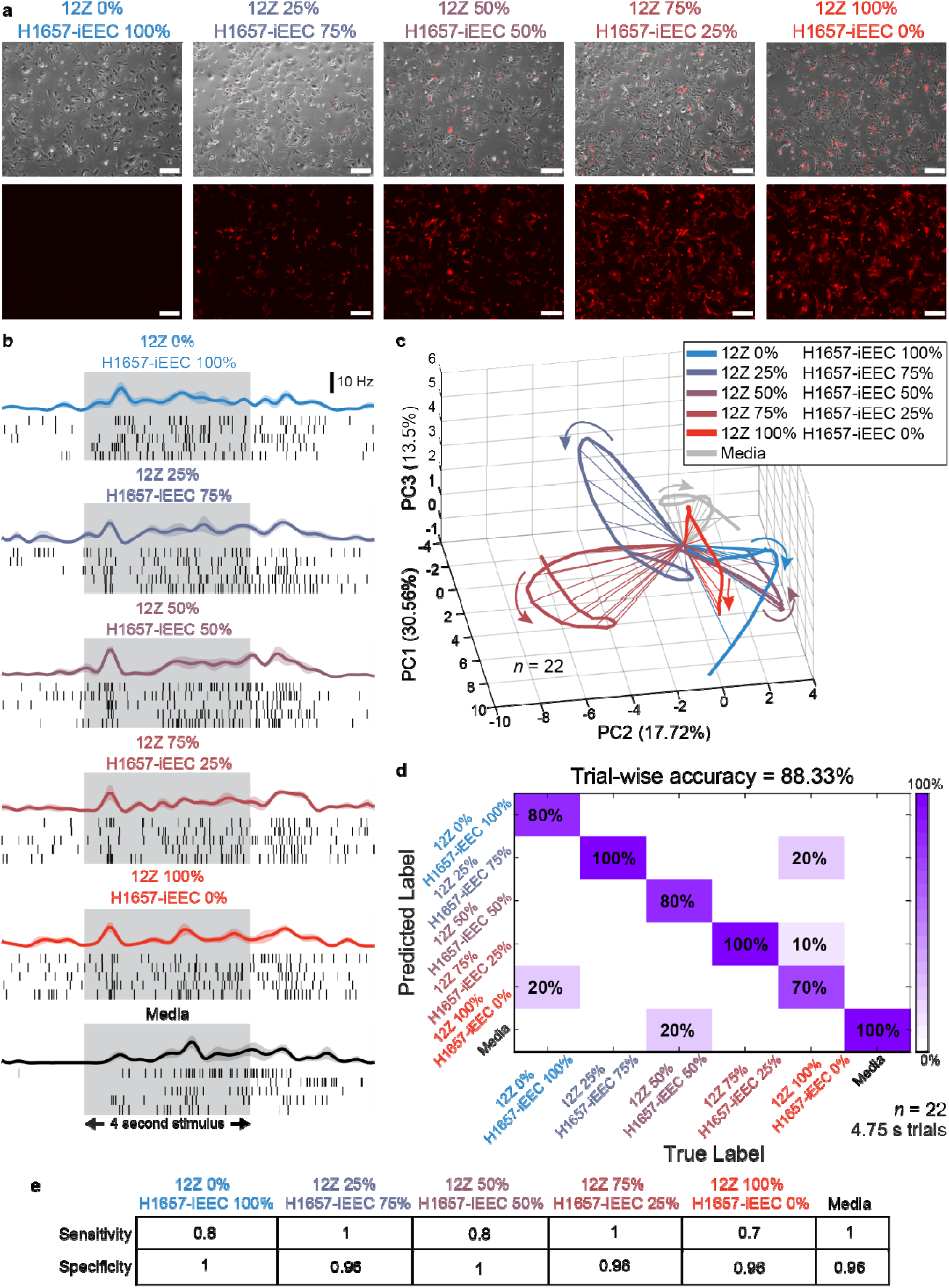
Differentiation and classification of co-cultures of endometriotic and endometrial cell lines. **a** Representative image of the epithelial cell co-culture combining endometriotic (12Z) and endometrial (H1657-iEEC) cell lines at multiple ratios. Immortalized human ectopic endometriotic epithelial cells (12Z) and immortalized endometrial epithelial cells (H1657-iEEC) are also shown without any co-culture. All the images are shown at 72-hour post-seeding. The white scale bar indicates 100 µm. **b** Distinct and tuned neural spiking responses to all cell culture, co-culture and media headspace gas mixtures is shown for one representative neuron. Representative raster plots show spiking patterns from five trials of each cell line and media. Each black line in the raster plot indicates an action potential, or spiking event, for this neuron. PSTHs are shown above each raster plot. Trial-averaged PSTHs are plotted with the shaded region indicating the SEM. The gray box indicates the 4 second stimulus presentation window. **c** Dimensionality reduction of high dimensional neural response vectors via PCA shows distinct temporal neural trajectories in the PCA subspace. Arrows specify the direction of the neural trajectories from the origin which indicates the start of the time window. The time window plotted is 0.5 – 1.75 seconds after stimulus onset. **d** Confusion matrix summarizing the high dimensional LOTO analysis for the classification of 4.75 second trials for the ‘on’ response time window of 0.25 – 2.25 seconds after stimulus onset in combination with the ‘off’ response time window of 3.25 – 6 seconds after stimulus onset (see Supplementary Fig. 11a). **e** Sensitivity and specificity table summarizing the results from the confusion matrix in panel d. Data was RMS filtered (see Methods) from 22 electrophysiological recording locations for analysis in panels c and d.

Employing a combined “on” and “off” response trial template (Supplementary Fig. 11a) for the high-dimensional LOTO analysis we obtained an accuracy of 88.33% for classifying all cell lines, co-cultures, and media (Fig. 5d) showcasing the robust ability of this sensor to not only to detect and classify separately grown cell cultures but also cell lines co-cultured together in varying progressive ratios. Remarkably, we also obtained a high classification accuracy of 95% using the population of spike sorted neurons with the high dimensional LOTO analysis (Supplementary Fig. 11b). The sensitivity and specificity of the sensor to classify each of the co-cultures, individual cell cultures and media was calculated summarizing the results from the LOTO analysis (Fig. 5e and Supplementary Fig. 11c). To validate the resulting classification accuracy, we performed a LOTO analysis on the 2 second baseline neural response recorded prior to stimulus onset and obtained an accuracy of 8.33% (Supplementary Fig. 12). Additional experiments were conducted using the stromal cell lines (iEc-ESC; endometriotic and H1644-iESC; endometrial), however the resulting analysis showed that these co-cultures were not classified with high classification accuracy (i.e., 51.67%) with some individual co-cultures classifying higher than others (Supplementary Fig. 13).

## Discussion

This study demonstrates that an indirect holistic measure of emitted volatile organic metabolites is sufficient to classify pathogenic vs control samples providing a systemic approach to endometriosis detection. We utilized a biological sensing and information encoding system along with computational analyses of neural data to differentiate multiple human endometriotic cell lines, offering a novel method of detection for a complex disease that does not rely on the identity or quantification of specific biomarkers. The investigation and proposal of candidate biomarkers across several biological compartments and fluids suggest the broad molecular modifications that result due to the processes of the disease ^23,24^. Taking a nonspecific multiplexed approach to endometriosis detection, we can utilize the many by-products of endometriosis associated metabolism, inflammation, and other processes bypassing the necessity for specific biomarker identification and quantification ^5^. Using four cell lines representing endometriotic and endometrial models grown in modified airtight cell culture flasks allowed us to deliver the contained mixture of volatile organic metabolites to the insect antenna while simultaneously recording from the insect antennal lobe. By doing this, we obtained single unit activity that responded significantly different to multiple stimulus conditions implying that the emitted gas mixtures were giving repeated distinct responses over the population of individually recorded neurons. To further probe the clinical applicability of our sensor we implemented an efficient unsupervised data processing technique to reduce the computational workload of the analysis while also keeping the discriminatory features of the neural response. Spike sorting neural data is the standard to obtain single units associated with the neural response and is an essential technique for understanding the neurobiological basis of behavior, neurological processes, and connectivity across different brain regions however also requires human supervision and has variability due to human performance ^75,76,84^. Processing neural data efficiently and unsupervised could allow for a point-of care diagnostic and here we show that RMS-based data processing technique is suitable for neural voltage signals evoked by endometriotic samples ^85^. By using both low and high dimensional population analyses we obtained both qualitative and quantitative measurements for the differentiation and classification of multiple cell lines. Using a LOTO cross validation method we obtained an accuracy of 92% in classifying all four human endometriotic and endometrial cell lines and media. Of note, this classification accuracy occurs when taking into account the discriminatory features of the whole trial in which the stimulus response was analyzed supporting the importance of the spatiotemporal neural response in differentiating odors from previous studies ^70,71,73^.

As endometriosis tends to show a progressive nature, it is essential to be able to differentiate the growth of lesions over time ^2,86^. At the population level using our LOTO cross validation method we obtained a classification accuracy of 89% with only three cell lines (24 hr. 12Z, 72 hr. H1657-iEEC, and 24 hr. H1644-iEEC) showing slight misclassifications. Of note, the 24 hr. 12Z and the 24 hr. H1644-iEEC were misclassified to their 72-hr. counterpart. Conducting our LOTO analysis on baseline neural activity showed below random classification validating that only odor-evoked neural responses are differentiable and contribute to the high classification accuracy. In addition, we can achieve a sufficient sensitivity and specificity in comparison to other proposed biomarkers ^16,23,24,26^. Our results imply a robust ability to detect not only cell lines but also growth timepoints of cells suggesting there is concentration difference in VOC mixtures between the same and different cell lines that is distinguishable by the sensor.

To further understand the limits of our sensor we co-cultured cell lines together combining endometriotic and endometrial cells within the same flasks at multiple ratios. Pathogenic and non-pathogenic epithelial cells co-cultured at 0%, 25%, 50%, 75%, and 100% in ascending/descending ratios were able to be classified using the LOTO cross validation method with an accuracy of 88% implying that we can also distinguish the subtle differences of gas mixtures pertaining to the progression of pathogenic related VOCs among the mixture.

Interestingly, while the co-cultures of the epithelial cells could be distinguished, we did not see the same results for the stromal cells complementing previous studies indicating that the epithelium is important for driving pathologic processes while endometriotic stromal cells from lesions show minimal transcriptome alterations and a reduced capacity for cellular differentiation and decidualization ^87,88^. Our results suggest that the emitted volatile metabolites from both pathogenic and normal stromal cells are similar and support the hypothesis that endometriotic lesions may originate from the endometrium ^2,89^. It is unclear if there may be potential cell-cell interactions that may influence the emitted volatile metabolites however monoculture and co-cultures of cancer cell VOCs have been studied previously with changes being noted ^90^. This work investigates two cell types (epithelial and stromal) but does not take into account the multiple subtypes that endometriosis is generally categorized into, peritoneal endometriosis, ovarian endometriosis, and deep infiltrating endometriosis ^91^. The endometriotic epithelial cell line 12Z is derived from human peritoneal endometriosis while the stromal cell line iEc-ESC is derived from a human ovarian endometriotic cyst indicating a lack of diversity that could be further explored using cell lines derived from other subtypes ^67,68,92^. Although a focus of future work should take into account the importance of detecting peritoneal endometriosis as this phenotype represents the most frequent occurring case and has been proven difficult to diagnosis from imaging modalities ^2,93,94^.

The clinical application of this work still requires further validation using human samples. While the study of VOCs associated with endometriosis is rare, the use of gas chromatography-mass spectrometry (GC-MS) has been used previously to study human patient serum samples finding several differences in metabolites between experimental and control groups ^95,96^. It should be noted that within these studies preprocessing of patient samples in necessary and the GC-MS methodology is considered to be expensive and time-consuming which may hinder direct application in routine diagnostics ^96,97^. Another study using human urine samples found significantly different concentrations of 14 proteins having roles in immune system activation, iron metabolism, cell apoptosis, and proteolysis supporting the broad pathological responses that occur with the disease ^98^. The analysis of urine-based VOCs for cancer have been previously studied with results showing significant differences from control samples encouraging the question of whether VOCs emitted from endometriotic urine samples could be differentiated with this insect-based sensor methodology ^29,99–101^. Animal models of endometriosis have been used and provide a logical next step in the validation of this sensor ^102–104^. This work gives credence to the hypothesis that mixtures of VOCs associated with endometriosis are discriminatory and leads to the question of whether more complex biological fluids (e.g., urine and breath) would also have these discriminating volatile mixtures. We hypothesize as complexity of samples increases the need for advanced classification methods using machine learning techniques would need to be implemented. Early detection and treatment of endometriosis is important as this can lessen the burden and severity of impact to quality of life ^105^. A shift from surgical diagnosis to clinical diagnosis is essential for the initiation of therapy as early as possible ^4^. Overall, it is imperative to further understand the limits of detection of this bioelectronic sensing system for the early detection of endometriosis deepening the potential translational and clinical impact.

## Methods

### Cell lines

All cell lines were cultured at 37°C 5% CO_2_. The immortalized endometriotic epithelial 12Z cells (a gift from Anna Starzinski-Powitz at Goethe University, Frankfurt, Germany) and 12Z expressing RFP (12Z RFP) were maintained in Dulbecco’s Modified Eagle Medium/Nutrient Mixture F-12 (DMEM/F12) (Gibco) supplemented with 10% charcoal stripped fetal bovine serum (Millipore Sigma), 1% penicillin/streptomycin (Gibco), and 0.1% sodium pyruvate (Gibco) ^67,83^. The immortalized endometrial stromal cells (H1644-iESC), established from normal endometrium in the Fazleabas lab, and the H1644-iESC expressing azurite blue, were cultured in DMEM/F12 media supplemented with 10% charcoal stripped fetal bovine serum, 1% penicillin/streptomycin, 0.1% sodium pyruvate, and 10ug/mL hygromycin B. The immortalized ectopic endometriotic stromal cells (iEc-ESC), established from an endometrioma in the Fazleabas lab, were cultured in DMEM/F12 media supplemented with 10% charcoal stripped fetal bovine serum, 1% penicillin/streptomycin, 0.1% sodium pyruvate, and 10ug/mL hygromycin B ^68^. The immortalized endometrial epithelial cells (H1657-iEEC) established from normal endometrium in the Fazleabas lab, were cultured in DMEM/F12 media supplemented with 2% charcoal stripped fetal bovine serum, 1% penicillin/streptomycin, and 0.1% sodium pyruvate. All cell lines were grown T75 flasks until ∼90-100% confluency.

### Single cell line culture preparation

The 12Z, H1657-iEEC, iEc-ESC, and H1644-iESC cell lines were plated in their own individual T25 flask (Nunc™ EasYFlask™ 156340, Thermo Fischer Scientific, MA, USA). Airtight T25 flasks were modified prior to plating by placing inlet and outlet 19-gauge needles into each individual flask and setting each needle using a low-volatile, two-part epoxy at least 24-hr before cell plating. All cell lines were trypsinized using 2x Trypsin-EDTA (0.5%) (Gibco) to lift cells from the bottom of the T75 flasks. Cells were counted using Invitrogen Countess II Cell Counter and plated at 3×10^5 cells/flask (6×10^4 cells/mL) in the T25 flask. The cells were cultured in 5mL DMEM/F12 media supplemented with 10% charcoal stripped fetal bovine serum, 1% penicillin/streptomycin, and 0.1% sodium pyruvate for 72 hrs. A T25 flask containing 5mL DMEM/F12 media supplemented with 10% charcoal stripped fetal bovine serum, 1% penicillin/streptomycin, and 0.1% sodium pyruvate and no cells was also prepared at the same time and maintained at the same time as the flasks containing cells. After either 72- or 24-hours cells were imaged and transported to the Saha lab for analysis. Live cell brightfield images were captured for each flask on the day of transportation using a Nikon Eclipse TS100 inverted microscope.

### Cell co-culture preparation

The 12Z RFP and H1657-iEEC were trypsinized using 2x Trypsin-EDTA (0.5%) to lift cells from the bottom of the T75 flasks. Cells were counted using Invitrogen Countess Cell II Counter. The cells were combined at 25%, 50%, 75% concentration with a total cell density at 3×10^5 cells/flask (6×10^4 cells/mL) in the T25 flask (Nunc™ EasYFlask™ 156340, Thermo Fischer Scientific, MA, USA). Airtight T25 flasks were modified prior to plating by placing inlet and outlet 19-gauge needles into each individual flask and setting each needle using a low-volatile, two-part epoxy at least 24-hr before cell plating. A flask containing only 12Z RFP cells and a flask containing only H1657-iEEC were prepared at the same time. The cells were cultured in 5mL DMEM/F12 media supplemented with 10% charcoal stripped fetal bovine serum, 1% penicillin/streptomycin, and 0.1% sodium pyruvate for 72 hours. A T25 flask containing 5mL DMEM/F12 media supplemented with 10% charcoal stripped fetal bovine serum, 1% penicillin/streptomycin, and 0.1% sodium pyruvate and no cells was also prepared at the same time and maintained at the same time as the flasks containing cells. After 72 hours cells were imaged and transported to the Saha lab for analysis. Live cell brightfield and fluorescent images were captured for each flask on the day of transportation using a Nikon Eclipse TS100 inverted microscope. The fluorescent and bright field images were merged using Nikon Elements BR version 5.42.03.

The iEc-ESC and H1644-iESC azurite blue were trypsinized using 2x Trypsin-EDTA (0.5%) to lift cells from the bottom of the T75 flasks. Cells were counted using Invitrogen Countess Cell II Counter. The cells were combined at 25%, 50%, 75% concentration with a total cell density at 3×10^5 cells/flask (6×10^4 cells/mL) in the T25 flask. A flask containing only iEcESC and a flask containing only H1644-iESC azurite blue were prepared at the same time. The cells were cultured in 5mL DMEM/F12 media supplemented with 10% charcoal stripped fetal bovine serum, 1% penicillin/streptomycin, and 0.1% sodium pyruvate for 72hrs. A T25 flask containing 5mL DMEM/F12 media supplemented with 10% charcoal stripped fetal bovine serum, 1% penicillin/streptomycin, and 0.1% sodium pyruvate and no cells was also prepared at the same time and maintained at the same time as the flasks containing cells. After 72 hours cells were imaged and transported to the Saha lab for analysis. Live cell brightfield and fluorescent images were captured for each flask on the day of transportation using a Nikon Eclipse TS100 inverted microscope. The fluorescent and bright field images were merged using Nikon Element BR version 5.42.03.

### Cell culture headspace delivery

A commercial olfactometer (Aurora Scientific, 220A) was used for precision cell culture headspace stimulus delivery (Fig. 1a). At the start of each set of trials, 200 standard cubic centimeters per minute (sccm) of zero contaminant air was passed through the air flow line via a 1/16 in. diameter PTFE flow line to the locust antenna demoted the stimulus flow line. The end of the stimulus flow line was placed approximately 2-3 cm from the most distal antennal segment. An additional 200 sccm of zero contaminant air was passed through a separate flow line to the exhaust denoted the dilution flow line. Five seconds before odor stimulus delivery, 40% (80 sccm) of the dilution flow line was redirected through the cell culture flow line directly upstream of the cell culture flasks. The dilution flow line and the cell culture flow line joined downstream of the cell culture flasks. This allowed for the complete mixing of the 80 sccm cell culture headspace flow with the 120 sccm of the dilution flow line’s clean air. The air-volatiles mixture primed the line with volatiles up to the final valve, where the combined cell culture headspace + dilution flow was delivered to the exhaust. Upon stimulus onset, the final valve redirected the clean air flow to the exhaust and the cell culture headspace + dilution flow to the locust antenna via the stimulus flow line. After 4 seconds of constant flow and stimulus delivery, the final valve redirected the clean air flow back to the locust antenna via the stimulus flow line, alleviating potential headspace gas depletion during the cell culture headspace delivery. This protocol was designed to keep a constant flow rate through the stimulus flow line, thereby eliminating any potentially confounding neuronal responses due to changes in air pressure via mechanosensory detection. A 6-inch diameter funnel pulling a slight vacuum was positioned immediately behind the locust the locust during cell culture headspace stimulus delivery to ensure rapid removal of volatiles. Each 4 second-duration stimulus was repeated 5 times for a total of 5 trials with an interstimulus interval of 60 seconds. The order of the stimuli was pseudorandomized for each experiment.

### Electrophysiology

All neural recordings were performed on post-fifth instar locust (Schistocerca americana) of either sex raised in a crowded colony. For in vivo extracellular neural recordings, locusts were immobilized on a surgical platform and antennae were stabilized. A batik wax bowl around the head was constructed to isolate the region and then filled with a room temperature, physiologically balanced locust saline solution. The brain was kept hydrated with locust physiological saline solution described by Laurent *et al*. (in mM as follows: 140 NaCl, 5 KCl, 5 CaCl2, 4 NaHCO3, 1 MgCl2, 6.3 HEPES, pH 7.0; all chemicals from Sigma-Aldrich) ^61,66^. The removal of the exoskeleton between the antennae was performed and glandular tissue was removed until the brain was fully visible. Treatment with protease was done to remove the thin membrane sheath on top of the antennal lobes. This protocol followed previously published methods ^56^. Following surgery, the animals were placed in a Faraday cage isolated bench and a silver-chloride ground wire was placed in the saline bath. A commercial Neuronexus 16-channel silicon probe (A2×2-tet-3mm-150-150-121) with impedances between 200 and 300 kΩ was inserted into the locust AL (Fig. 1a) for all neural recordings. To complete the circuit via the saline solution, a silver chloride ground wire was placed within the head of the locust. Voltage signals were sampled at 20 kHz and then digitized using an Intan pre-amplifier board (C3334 RHD 32-channel head stage). The digitized signals were transmitted to the Intan recording controller (C3100 RHD USB interface board) before being visualized and stored using the Intan graphical user interface and LabView data acquisition system. Each neural recording lasted 1-2 hr.

### Spike sorting

All neural data was imported into MATLAB after high pass filtering using a 300Hz Butterworth filter to eliminate frequencies below 300 Hz. The data was analyzed by custom-written codes in MATLAB R2023b. All data was processed with Igor Pro for spike sorting analysis using previously described methods ^69^. Spiking events were identified using a detection threshold between 2.5 and 3.5 standard deviation (SD) of baseline fluctuations. Individual projection neurons were identified if they passed the following criteria: cluster separation > 5 SD, inter-spike intervals (ISI) ≤ 10%, and spike waveform variance ≤ 10%.

### Scatter plots

The total number of spikes for each spike sorted neuron throughout the 4 second stimulus presentation was calculated for each trial (5 trials total for each stimulus). The mean spike counts ± SEM across the trials for each neuron were subsequently plotted for two stimulus conditions along the X- and Y-axis. One-way ANOVA with Bonferroni correction due to multiple comparisons was utilized to determine if each neuron had statistically significant differences in mean spike counts to different stimulus conditions (P < 0.05, d.f. = 4, 20; single time point and d.f. = 8, 36; multiple timepoint experiments, one-way ANOVA with Bonferroni correction). Neurons that showed a significant increase or decrease in spike counts along the stimulus on the Y-axis in comparisons to the stimulus on the X-axis were plotted in red or blue, respectively. Neurons that showed statistically nonsignificant differences were plotted in grey.

### Root mean squared (RMS) transformation

All neural data was imported into MATLAB after high pass filtering using a 300Hz Butterworth filter to eliminate frequencies below 300 Hz. Large and broad voltage peaks, caused by electrical interference or animal movement that were greater than 15 standard deviations from the mean voltage amplitude were removed by setting the 200 samples centered around each peak equal to the mean voltage value. This was done to remove any artifact from the data. The filtered data was trimmed to the time window of interest and all data were passed through a 500-point continuous moving RMS filter followed by a smoothing step via a 500-point continuous moving average filter as described previously ^56,57^. Stimulus-specific baseline values were calculated as the average voltage over all time bins for the 2 seconds prior to the stimulus onset. Baseline responses were averaged over all trials and subsequently subtracted from the data to obtain the change in root mean squared (Δ RMS) values. These values were binned into non-overlapping 50 millisecond bins and the average of each bin was computed. For each recording location, RMS transformed voltage data of each 4-channel tetrode were averaged together.

### Dimensionality reduction analyses

We executed two techniques of dimensionality reduction – Principal Component Analysis (PCA) and Linear Discriminant Analysis (LDA) as described in our previous studies ^56,57^. In PCA, we binned baseline subtracted spike sorted or RMS transformed neural signals into 50 millisecond non-overlapping time bins and averaged across trials (n = 5; each cell line and media was repeated 5 times with a 60 second inter-stimulus interval). Stimulus-specific baseline values were calculated as the average voltage (RMS) or firing rate (spike sorted) over all time bins for the 2 seconds prior to the stimulus onset. The neural responses were pooled across electrophysiological experiments or spike sorted neurons to generate a matrix, where each element in the matrix corresponds to the neural responses of one location or neuron in one 50 millisecond time bin. Similar neural population time-series data matrices were generated for each cell line and media stimulus. PCA dimensionality reduction analysis was performed on the time-series data involving all cell lines including media control and directions of maximum variance were found. The consequent high-dimensional vector in each time bin was projected along the eigenvectors of the covariance matrix. Only the three dimensions with the highest eigenvalues were used for visualization and data points in subsequent time bins were connected to produce low-dimensional neural trajectories. The trajectories were smoothed using a third order IIR Butterworth filter (Half Power Frequency = 0.15). Lastly, all trajectories were shifted to begin at the origin to analyze stimulus-specific response dynamics and trajectory divergence (e.g., Fig. 2A). For LDA, the same neural population time-series data matrix was used. Here, we maximized the separation between interclass distances while minimizing the within class distances. For visualization purposes, time bins were plotted as unique points in this transformed LDA space and stimulus-specific clusters became noticeable (e.g., Fig. 2b). All dimensionality reduction analyses were accomplished using custom written MATLAB (R2023b) codes.

### Hierarchical clustering

Spike-sorted neural responses were binned into 50 ms non-overlapping time bins for hierarchical clustering analysis. Average baseline spiking responses were computed using the 2 seconds prior to stimulus onset and then subtracted throughout the dataset. The pairwise distance between each 50 ms time bin and the observations (e.g., 5 trials x 9 cell line panel = 45 observations) was computed using high dimensional (e.g., 58 neurons yield 58 dimensions) Euclidean distance. The distances were then averaged for each pairwise observation to output a single distance value. A hierarchical cluster tree was created through unsupervised agglomerative hierarchical clustering (‘linkage’ function in MATLAB) using the Ward agglomeration method. We performed hierarchical clustering analysis using custom written MATLAB (R2023b) codes.

### Quantitative classification analysis: leave-one-trial-out

To achieve a quantitative estimate of classification performance, we performed a leave-one-trial-out (LOTO) analysis. During each iteration, population neural response time-series data from one trial was used as the test data and the remaining four trials were used to train a linear classifier. This was repeated for all cell lines and media in the stimulus panel (e.g., for 4 cell lines and media there will be 5 training and 5 testing templates). The linear algorithm generated a model based on the training data set within the original high dimensional encoding state space to effectively classify testing data. By considering time bins as points in a high-dimensional space, we calculated an average response vector for the training data corresponding to each stimulus.

These neural templates were then used to classify individual time bins (50 millisecond duration) of each testing data set. The number of time bins depends on the time window if interest. The minimal Euclidean or Manhattan distance between each point corresponding to a time bin of the testing data and the previously calculated average response vectors for the training data were used to assign class identities. The Euclidean (L^2^) norm was used to quantify classifier predictability. We repeated this process leaving out a different trial to serve as the testing data and the other four trials to serve as the training data with each iteration. This type of analysis is termed a bin-wise classification (e.g., Fig. 2c). Furthermore, a winner-take-all approach, was also incorporated to calculate the most likely predicted class for each trial (e.g., Fig. 2d). This was performed by considering the mode of all predicted time bins as the trial-wise class identifier.

Model performance was illustrated using a confusion matrix, which compared the predicted responses to the true class labels. For example, a fully diagonal matrix indicates 100% classification accuracy. For a schematic representation of this analysis see our previous publication ^57^. We performed quantitative classification analysis using custom written MATLAB (R2023b) codes.

### Sensitivity and specificity

For the cell culture experiments at multiple timepoints, sensitivity and specificity were calculated with respect to each individual cell line and media. This is a non-binary classification as a trial can be classified as one of the 8 cell lines or media. For non-binary classifications where there are more than two possible outcomes, we used a one-vs-rest method to convert into a binary classification. Briefly, the odor of interest is defined as the positive odor and all other odors are combined and defined as the negative odor. This was repeated for all cell lines and media at separate timepoints. For calculating the overall sensitivity and specificity of a cell line we combined the 24 hr. and 72 hr. results. For example, in calculating the sensitivity and specificity of the 12Z cell culture, 24 hr. 12Z and 72 hr. 12Z were combined. For calculating the endometriotic sensitivity and specificity, all endometriotic cell lines (12Z and iEc-ESC) at all timepoints (24 hr. and 72 hr.) were combined. This was repeated for all endometrial cell culture excluding media. For the co-culture experimental analysis, sensitivity and specificity was calculated with respect to each individual cell line, co-culture, and media using the one-vs-rest method.

### Statistical analysis

Statistical analysis used one-way ANOVA with Bonferoni correction for multiple comparisons. Data are expressed as mean ± SEM, with P < 0.05 considered statistically significant.

## Data availability

Data analysis codes are available from the corresponding author on request. Data is available on Dryad.

## Acknowledgments

This work was supported by National Science Foundation Graduate Research Fellowship (2235783) to S.W.S., National Science Foundation CAREER Award (2238686) to D.S., National Institutes of Health grant (R01HD099090) to A.T.F, National Institutes of Health grant (1R21HD114955) to A.T.F. and D.S.

## Author contributions

Conceptualization: A.T.F., D.S. Methodology: S.W.S., E.L.V., Y.S., M.P., A.T.F., D.S. Investigation: S.W.S., E.L.V., Y.S., M.P., A.T.F., D.S. Visualization: S.W.S. Funding acquisition: S.W.S., A.T.F., D.S. Project administration: S.W.S., E.L.V., A.T.F., D.S. Supervision: S.W.S., E.L.V., A.T.F., D.S. Writing – original draft: S.W.S., E.L.V., A.T.F. Writing – review & editing: S.W.S., E.L.V., Y.S., M.P., A.F., D.S.

## Competing interests

Authors declare that they have no competing interests.

## Notes

### Competing Interest Statement

The authors have declared no competing interest.

