## Supplemental Data and Figures for "An insect brain-based bioelectronic neural sensor for the systemic detection and precise classification of endometriosis"

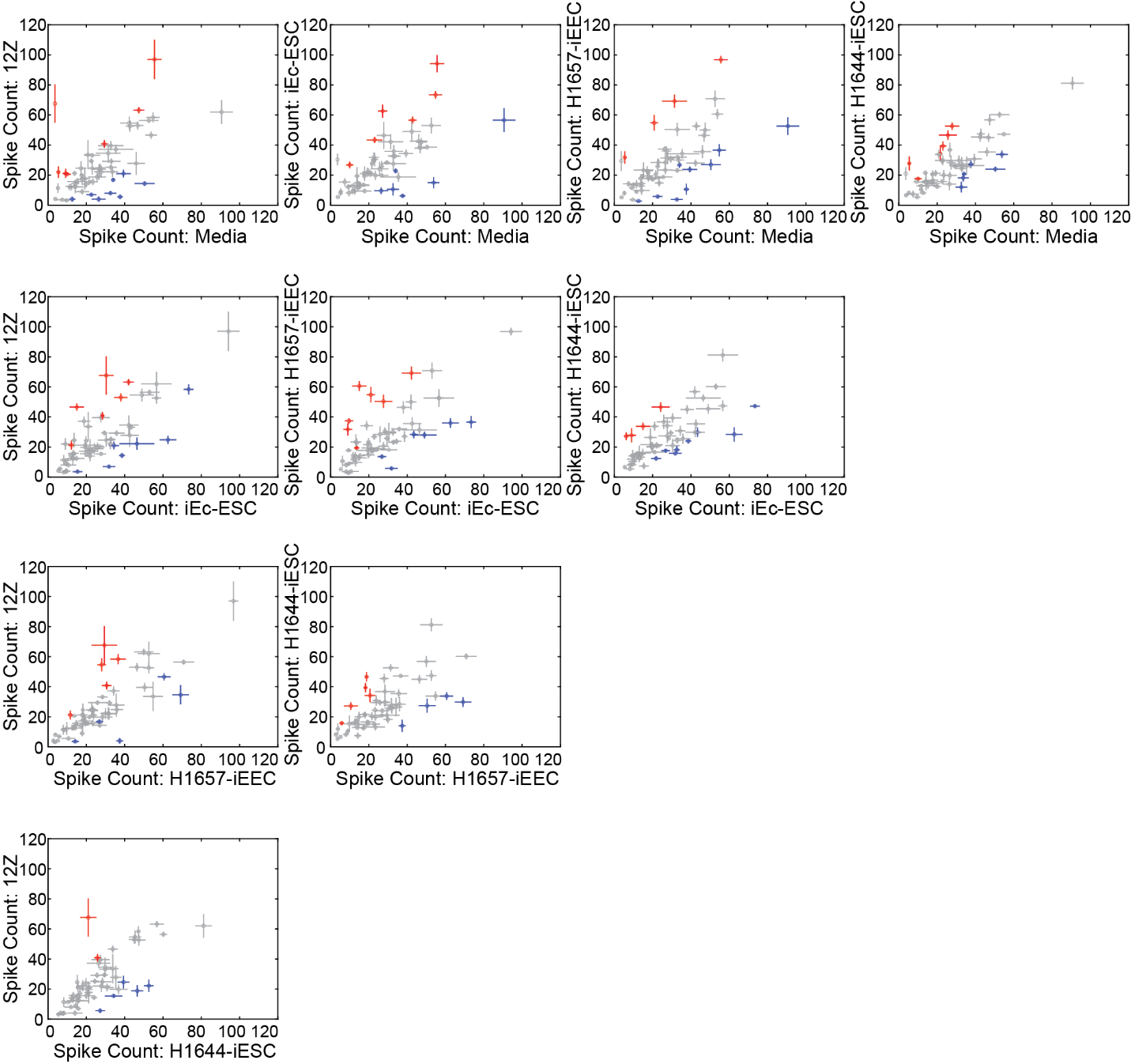

**Supplementary Figure 1.** **Neural spike counts show significant difference between two stimulus conditions.** Comparison of stimulus-evoked total spike counts (over 4 seconds) for 58 individual neurons spike sorted from all extracellular electrophysiological recordings for two stimulus conditions. Trial averaged total spike counts are plotted with error bars representing the standard error of the mean (SEM) of the trial-wise variations for two cell lines or media. Spike sorted neurons that responded significantly higher or lower compared to the x-axis stimulus were plotted in red or blue, respectively (p < 0.05, d.f. = 4, 20, one-way ANOVA with Bonferroni correction). Spike sorted neurons that did not show significant differences in total spike counts across the two conditions were plotted in grey.

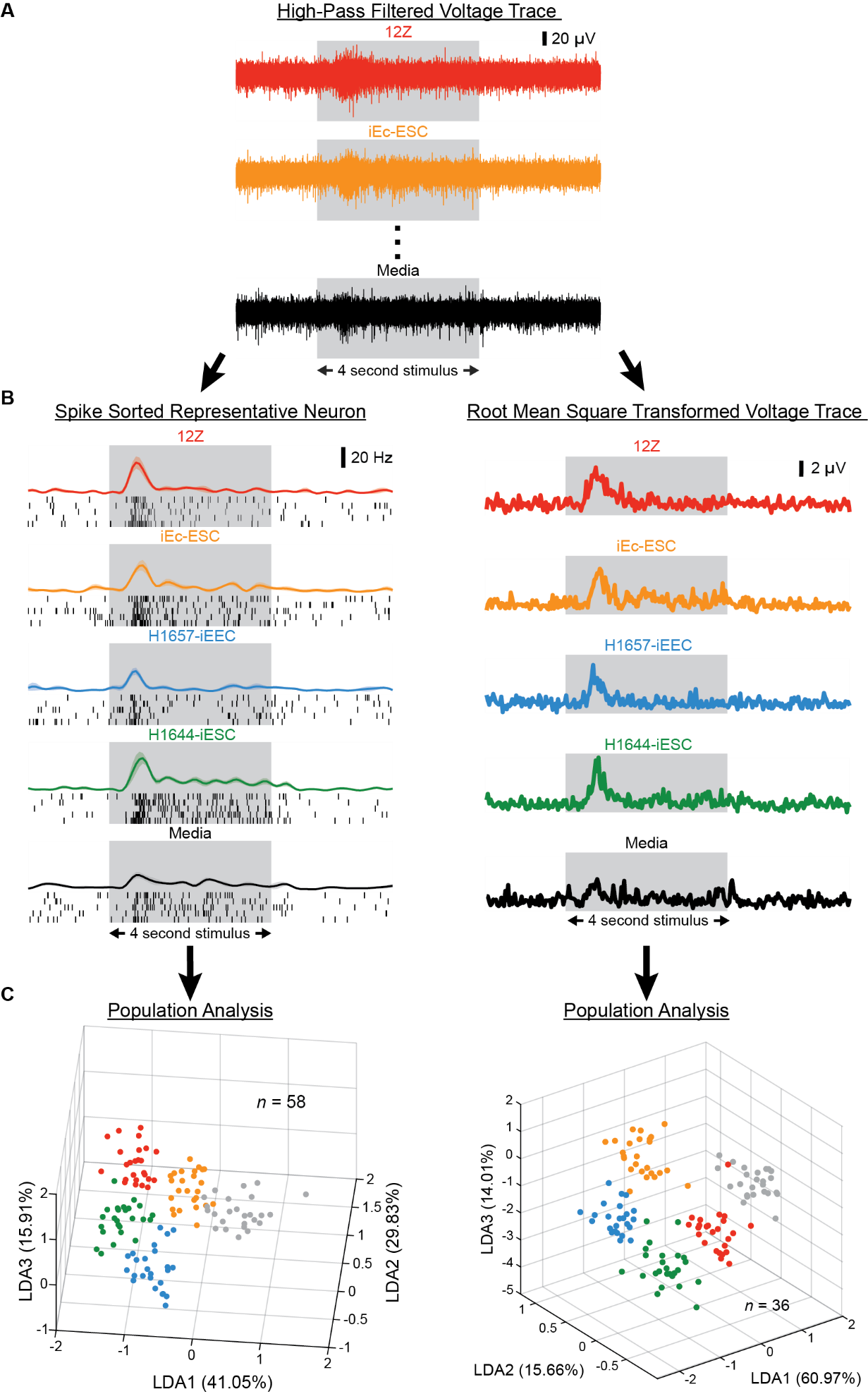

**Supplementary Figure 2. Schematic of parallel analysis workflow.** (**A**) Representative extracellular neural voltage responses obtained from the locust antennal lobe for each cell culture in the experimental panel. The grey box indicates the 4 second stimulus presentation window. (**B**) After the voltage response are obtained, the analysis workflow diverts into two parallel processing techniques: spike sorting or root mean squared (RMS) filtering (see Table S1. for a comparison of each neural processing technique). Representative output examples of each technique are shown. The grey box indicates the 4 second stimulus presentation window. (**C**) After processing the neural voltage responses, the output of each technique is included into an accumulative dataset for population analysis. A representative example of linear discriminant analysis using the dataset from each processing technique obtained is shown. In the spike sorted example, 58 neurons are obtained via spike sorting (see Methods) and used to for linear discriminant analysis (LDA). In the RMS example, 36 electrophysiological recording locations are filtered (see Methods) and used to for LDA. Each point in the LDA subspace indicates a 50-millisecond time bin.

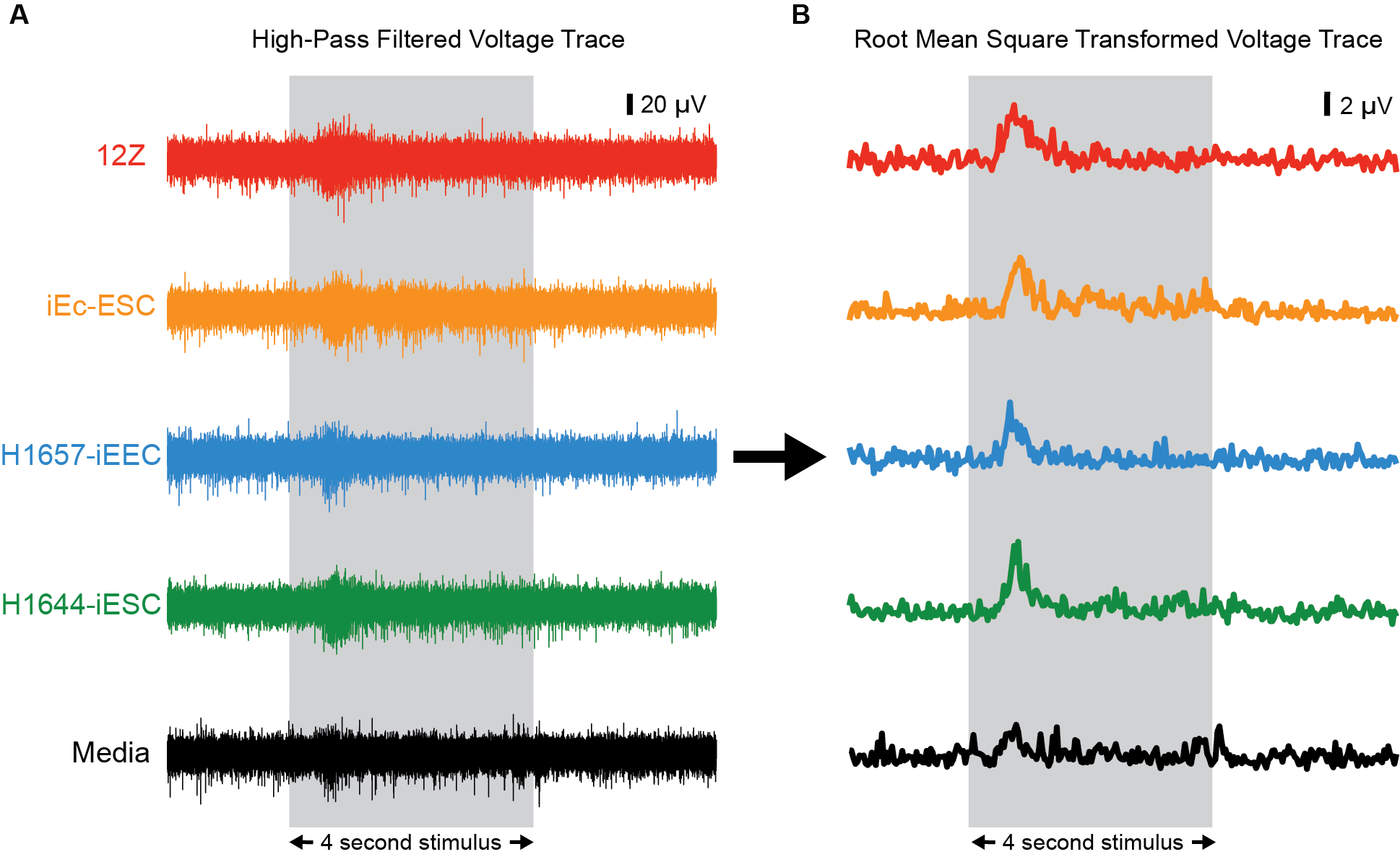

**Supplementary Figure 3. Root mean squared transformation preserved stimulus-specific spiking dynamics.** (**A**) Representative extracellular neural voltage responses obtained from an individual electrode are shown for all cell culture after high-pass filtering. The grey box indicates the 4 second stimulus presentation window. (**B**) RMS transformed data traces of (A) (see Methods) reflect the spiking rate-based response dynamics, while reducing the computational load for subsequent analysis. The grey box indicates the same 4 second stimulus presentation window in (A).

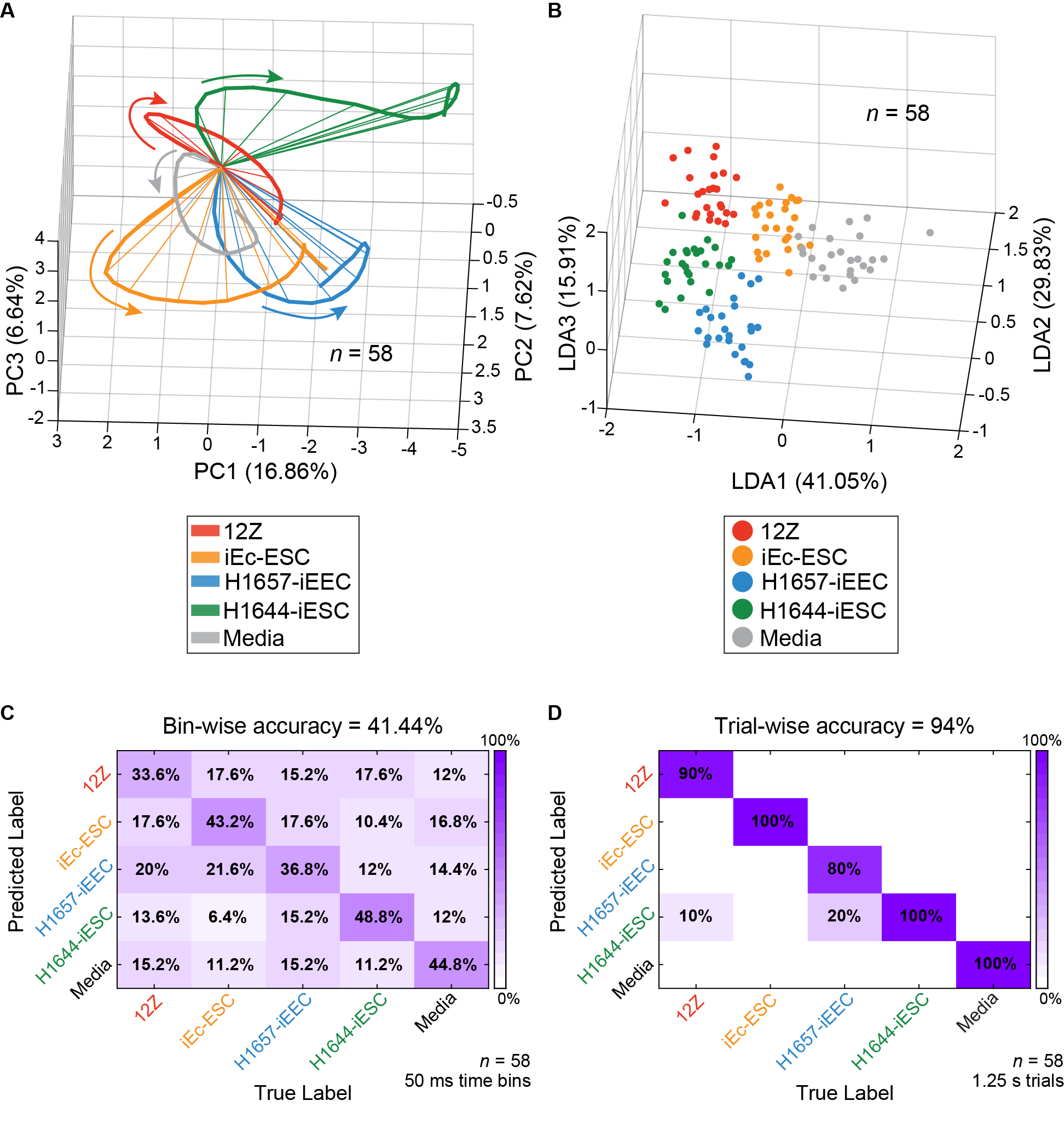

**Supplementary Figure 4. Spatiotemporal population neural responses distinguish and classify emitted endometriotic and healthy cell culture gas mixtures.** (**A**) Dimensionality reduction of high dimensional neural response vectors via principal component analysis (PCA) shows distinct temporal neural trajectories in the PCA subspace. Arrows specify the direction of the neural trajectories from the origin which indicates the start of the time window. The time window plotted is 0.5 – 1.75 seconds after stimulus onset. (**B**) Dimensionality reduction of high dimensional neural response vectors via LDA shows distinct neural response clusters in the LDA subspace. Each point in the LDA subspace indicates a 50-millisecond time bin within the time window of 0.5 – 1.75 seconds after stimulus onset. (**C**) Confusion matrix summarizing the high dimensional leave-one-trial-out (LOTO) analysis for the classification of 50 millisecond time bins within the time window of 0.5 – 1.75 seconds after stimulus onset. (**D**) Confusion matrix summarizing the high dimensional LOTO analysis for the classification of 1.25 second trials for the entire time window of 0.5 – 1.75 seconds after stimulus onset. 58 neurons were spike sorted (see Methods) from 36 electrophysiological recording locations for analysis.

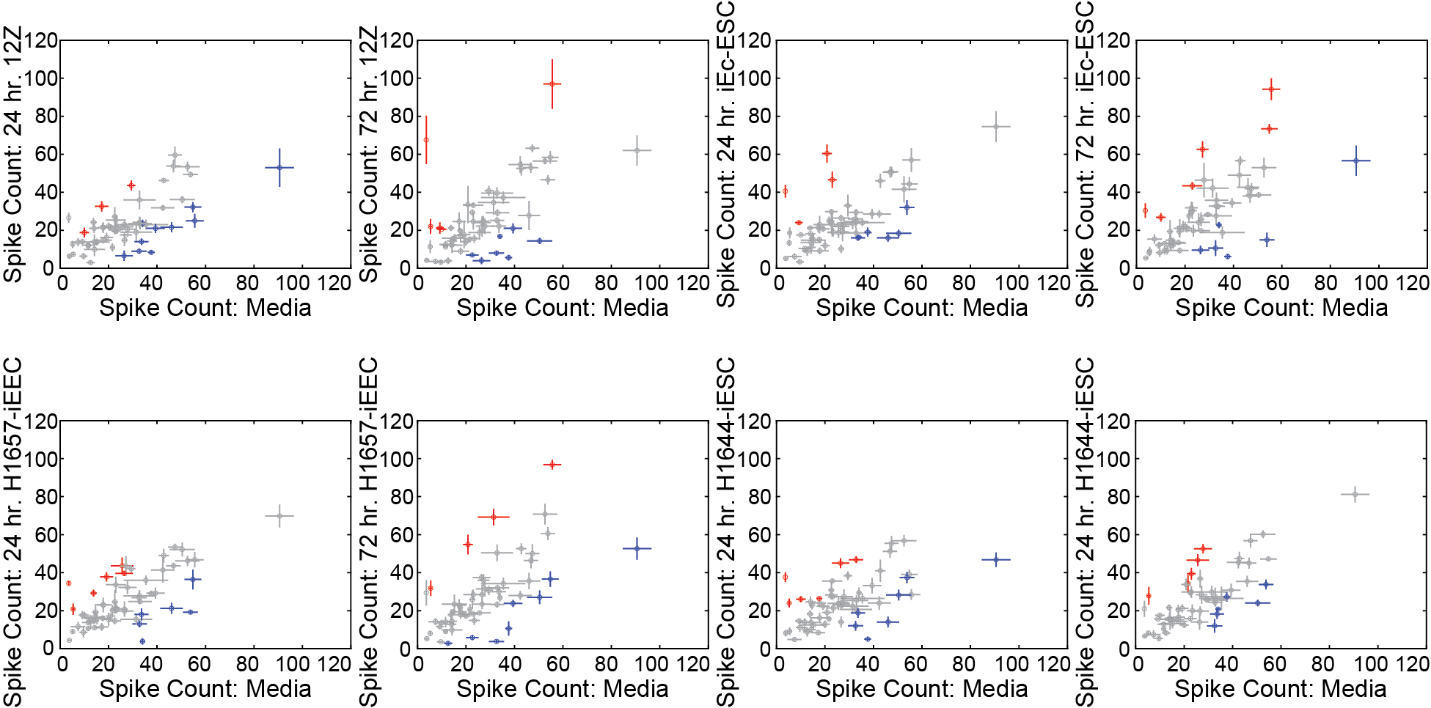

**Supplementary Figure 5. Neural spike counts show significant difference between two stimulus conditions for cell lines at multiple time points.** Comparison of stimulus-evoked total spike counts (over 4 seconds) for 58 individual neurons spike sorted from all extracellular electrophysiological recordings for two stimulus conditions. Trial averaged total spike counts are plotted with error bars representing the SEM of the trial-wise variations for two cell lines or media. Spike sorted neurons that responded significantly higher or lower compared to the x-axis stimulus were plotted in red or blue, respectively (p < 0.05, d.f. = 8, 36, one-way ANOVA with Bonferroni correction). Spike sorted neurons that did not show significant differences in total spike counts across the two conditions were plotted in grey.

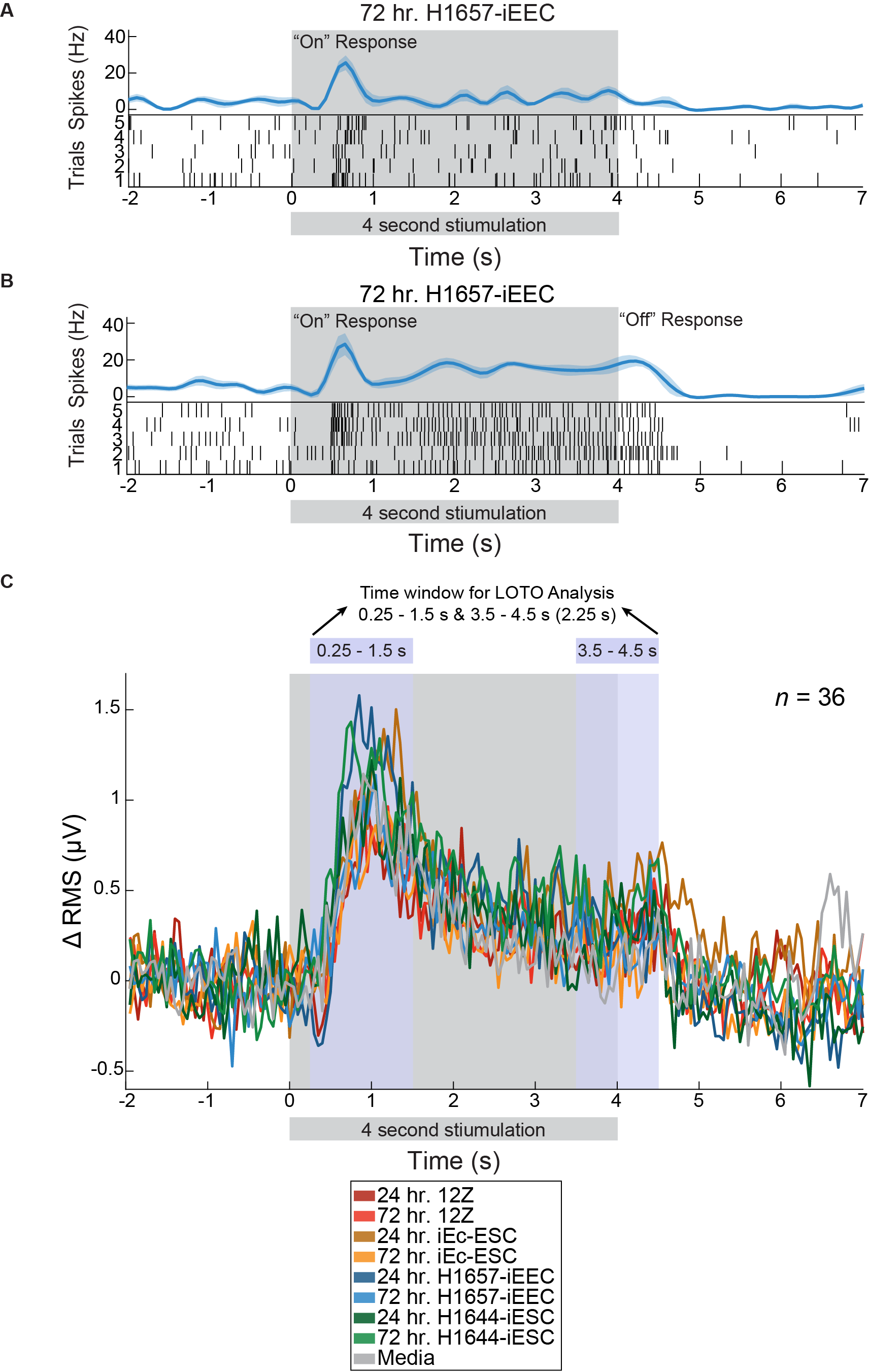

**Supplementary Figure 6. Neural “on” and “off” responses from cell culture emitted gas mixtures.** (**A**) Representative raster and PSTH for a single neuron’s response to the emitted gas mixture from 72 hr. H1657-iEEC are shown displaying an “on” response during the stimulus presentation window. Raster plots contain 5 trials, in which the gas mixture was delivered to the locust antennae. Each line in the raster plot indicates a spiking event, or action potential. PSTHs show the average changes in spiking rate for the 5 trials shown in the corresponding raster plot. Trial-averaged PSTHs are plotted with the shaded region indicating the SEM. The grey box indicates the 4 second stimulus presentation window. Time labels along the X-axis are relative to the stimulus onset time. (**B**) Representative raster and PSTH for a single neuron’s response to the emitted gas mixture from 72 hr. H1657-iEEC are shown displaying an “on” response during the stimulus presentation window and an “off” response after the stimulus presentation window. Raster plots contain 5 trials, in which the gas mixture was delivered to the locust antennae. Each line in the raster plot indicates a spiking event, or action potential. PSTHs show the average changes in spiking rate for the 5 trials shown in the corresponding raster plot. Trial-averaged PSTHs are plotted with the shaded region indicating the SEM. The grey box indicates the 4 second stimulus presentation window. Time labels along the X-axis are relative to the stimulus onset time. (**C**) Population based peri-stimulus time histogram is shown for all 36 electrophysiological recordings after RMS processing for each cell line. Time labels along the X-axis are relative to the stimulus onset time. The grey box indicates the 4 second stimulus presentation window. The blue boxes indicate the time window durations (encompassing the “on” and “off” response) used for LOTO analysis shown in Fig. 4D and fig. S8C.

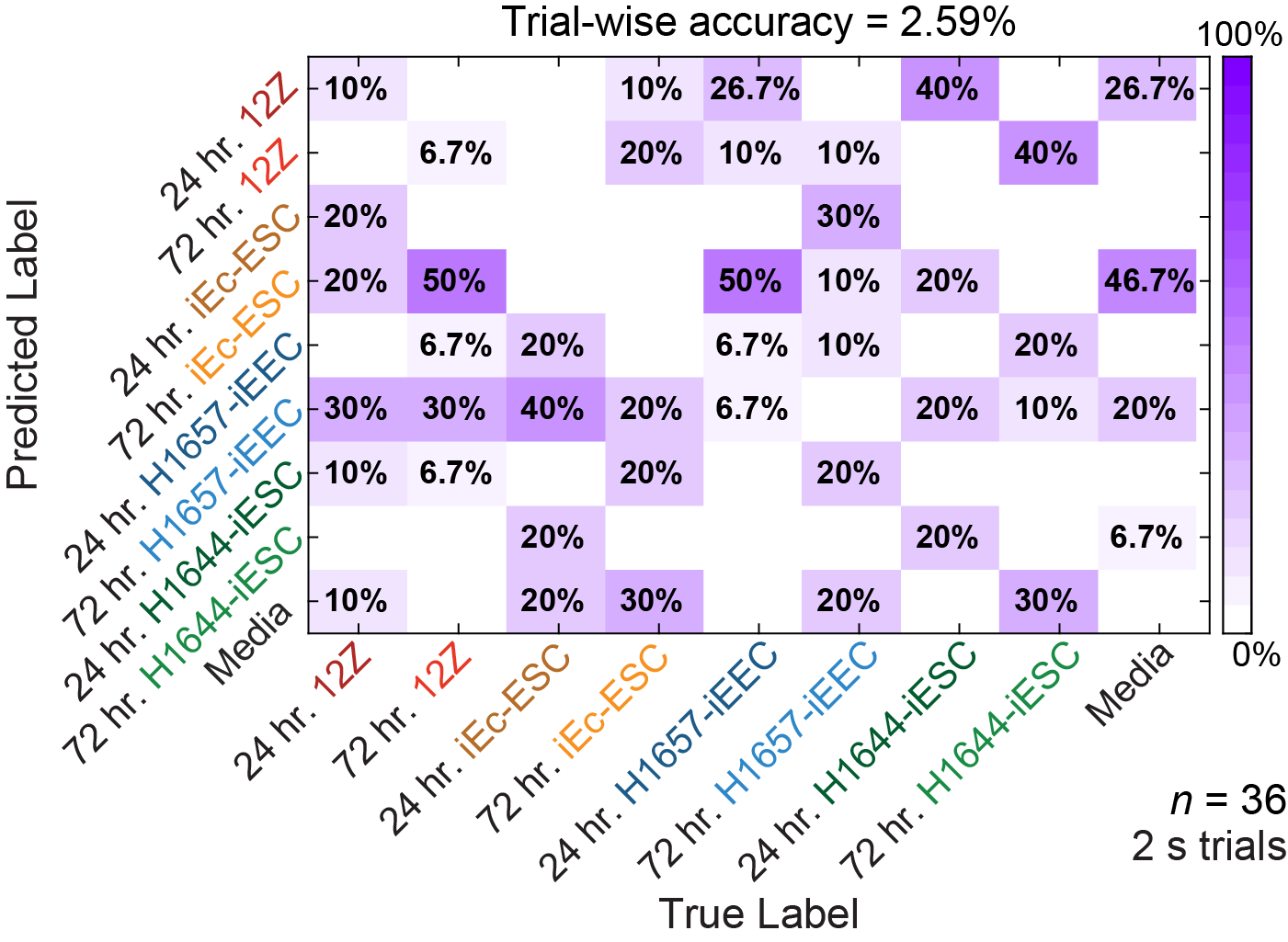

**Supplementary Figure 7. Confusion matrix summarizing the LOTO analysis for the two seconds prior to stimulus onset.** Confusion matrix summarizing the high dimensional LOTO analysis for the classification of 2 second trials for before the stimulus onset. Data was RMS filtered (see Methods) from 36 electrophysiological recording locations for analysis.

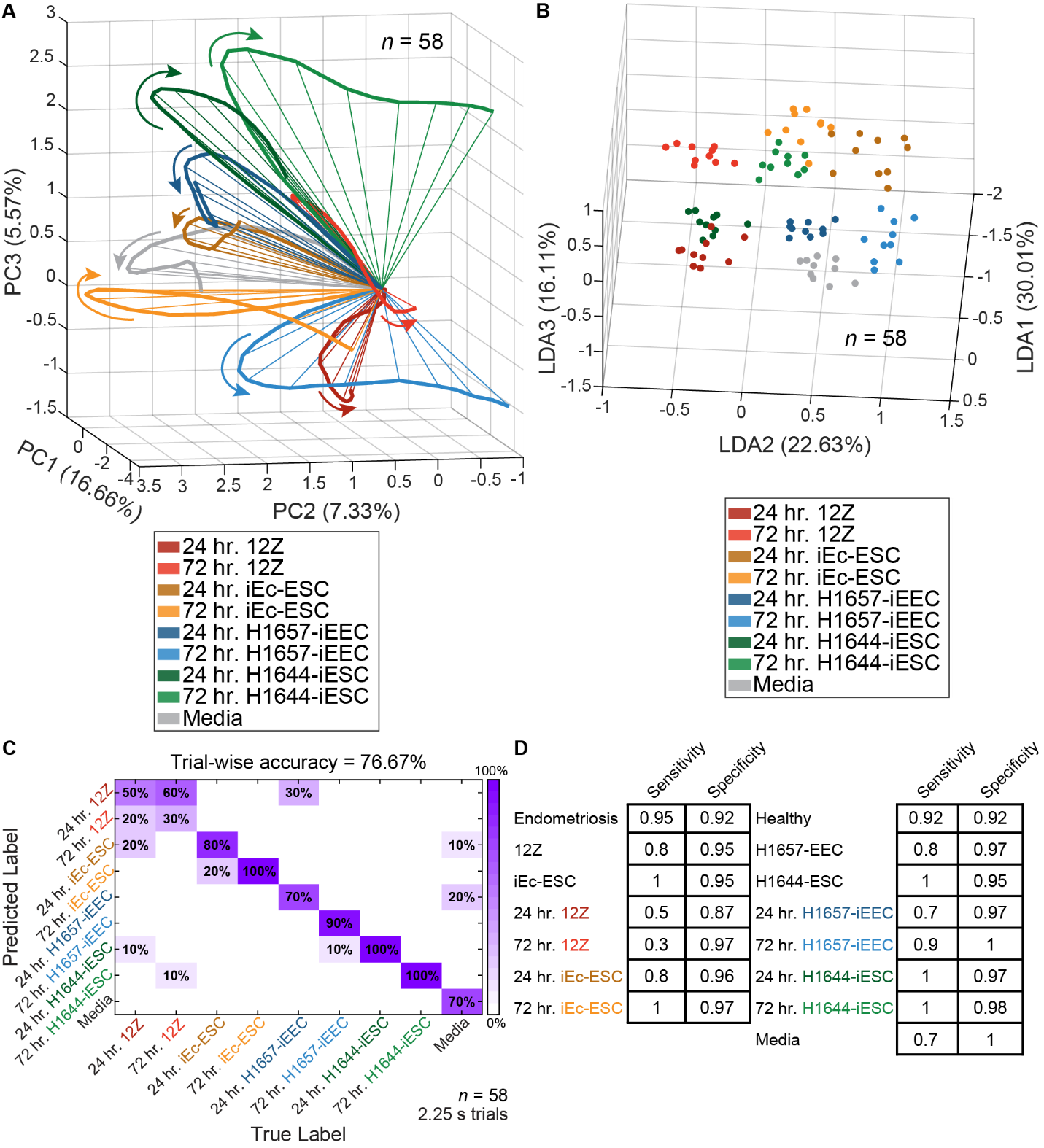

**Supplementary Figure 8. Spatiotemporal population neural responses distinguish and classify emitted endometriotic and endometrial cell culture gas mixtures at multiple time points.** (**A**) Dimensionality reduction of high dimensional neural response vectors via PCA shows distinct temporal neural trajectories in the PCA subspace. Arrows specify the direction of the neural trajectories from the origin which indicates the start of the time window. The time window plotted is 0.5 – 1.5 seconds after stimulus onset. (**B**) Dimensionality reduction of high dimensional neural response vectors via LDA shows distinct neural response clusters in the LDA subspace. Each point in the LDA subspace indicates a 50-millisecond time bin within the time window of 0.5 – 1 seconds after stimulus onset. (**C**) Confusion matrix summarizing the high dimensional LOTO analysis for the classification of 2.25 second trials for the ‘on’ response time window of 0.25 – 1.5 seconds after stimulus onset in combination with the ‘off’ response time window of 3.5 – 4.5 seconds after stimulus onset (see fig. S6). (**D**) Sensitivity and specificity table summarizing the results from the confusion matrix in (C). 58 neurons were spike sorted (see Methods) from 36 electrophysiological recording locations for analysis.

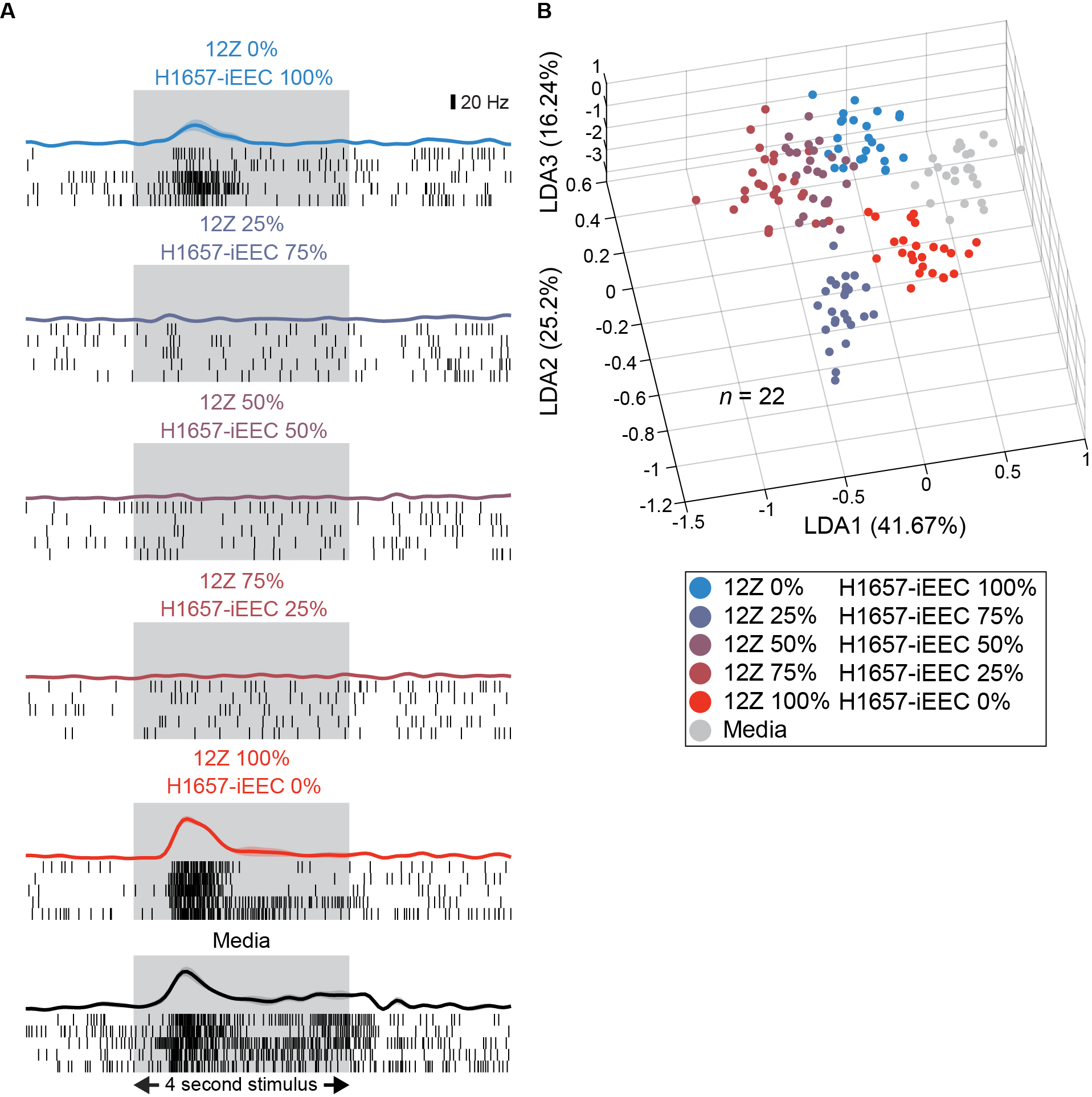

**Supplementary Figure 9. Individual and population neural responses differentiate cell co-culture emitted gas mixtures.** (**A**) Distinct neural spiking responses to cell culture, co-culture and media headspace gas mixtures is shown for one representative neuron. Representative raster plots show spiking patterns from five trials of each cell line and media. Each black line in the raster plot indicates an action potential, or spiking event, for this neuron. PSTHs are shown above each raster plot. Trial-averaged PSTHs are plotted with the shaded region indicating the SEM. The gray box indicates the 4 second stimulus presentation window. (**B**) Dimensionality reduction of high dimensional neural response vectors via LDA shows distinct neural response clusters in the LDA subspace. Each point in the LDA subspace indicates a 50-millisecond time bin within the time window of 0.5 – 1.75 seconds after stimulus onset. Data was RMS filtered (see Methods) from 22 electrophysiological recording locations for analysis in (B).

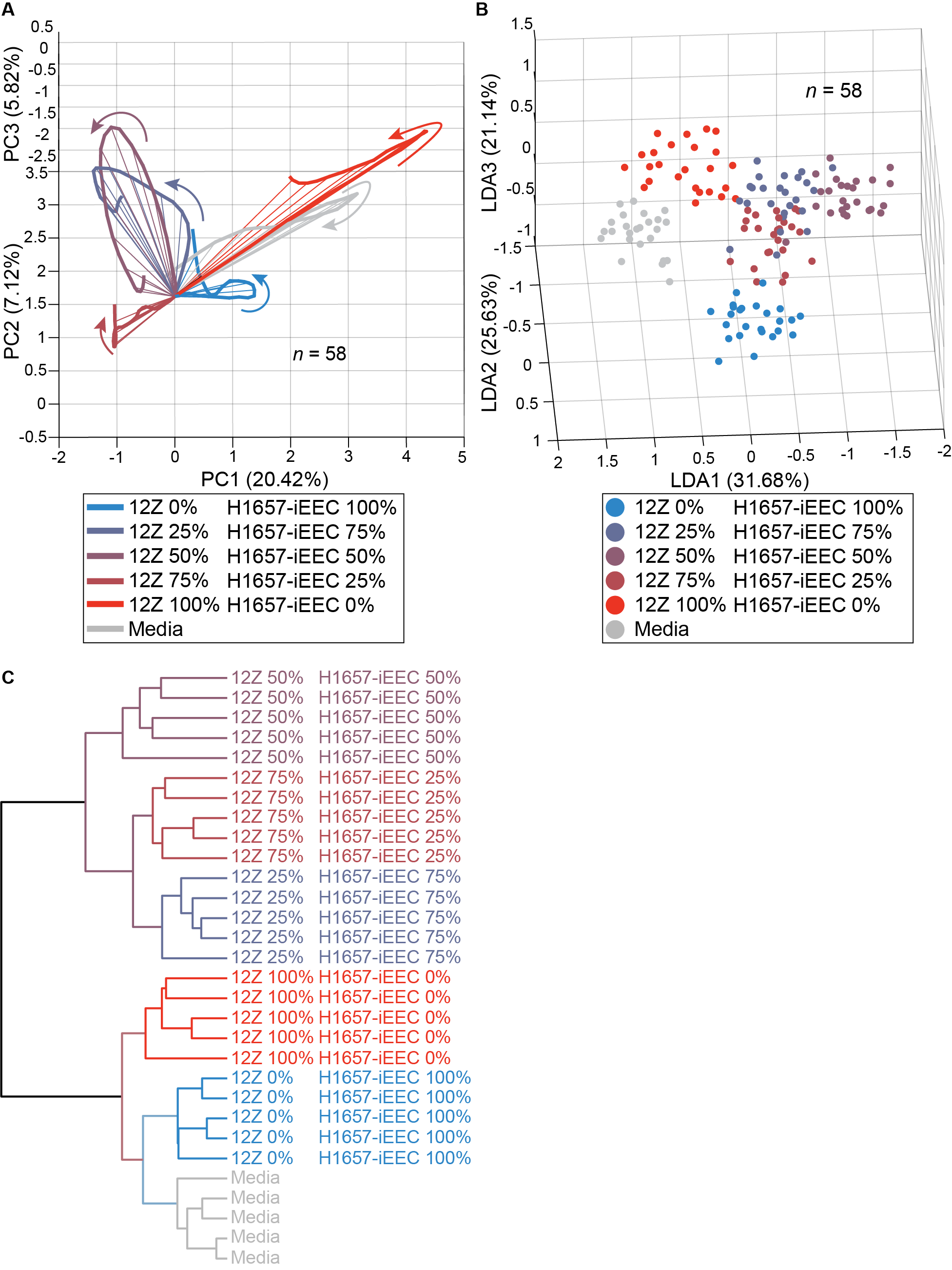

**Supplementary Figure 10. Spatiotemporal population neural responses distinguish emitted endometriotic and healthy cell co-culture gas mixtures at multiple ratios.** (**A**) Dimensionality reduction of high dimensional neural response vectors via PCA shows distinct temporal neural trajectories in the PCA subspace. Arrows specify the direction of the neural trajectories from the origin which indicates the start of the time window. The time window plotted is 0.5 – 1.75 seconds after stimulus onset. (**B**) Dimensionality reduction of high dimensional neural response vectors via LDA shows distinct neural response clusters in the LDA subspace. Each point in the LDA subspace indicates a 50-millisecond time bin within the time window of 0.5 – 1.75 seconds after stimulus onset. (**C**) A hierarchical clustering analysis dendrogram (see Methods) grouped the responses into two clusters separated by co-culture and individual cell lines. Moreover, these clusters were further grouped by the specific percentage of each cell line within the flasks. The time window for this analysis was 0.5 – 4 seconds after the stimulus onset. 58 neurons were spike sorted (see Methods) from 22 electrophysiological recording locations for analysis.

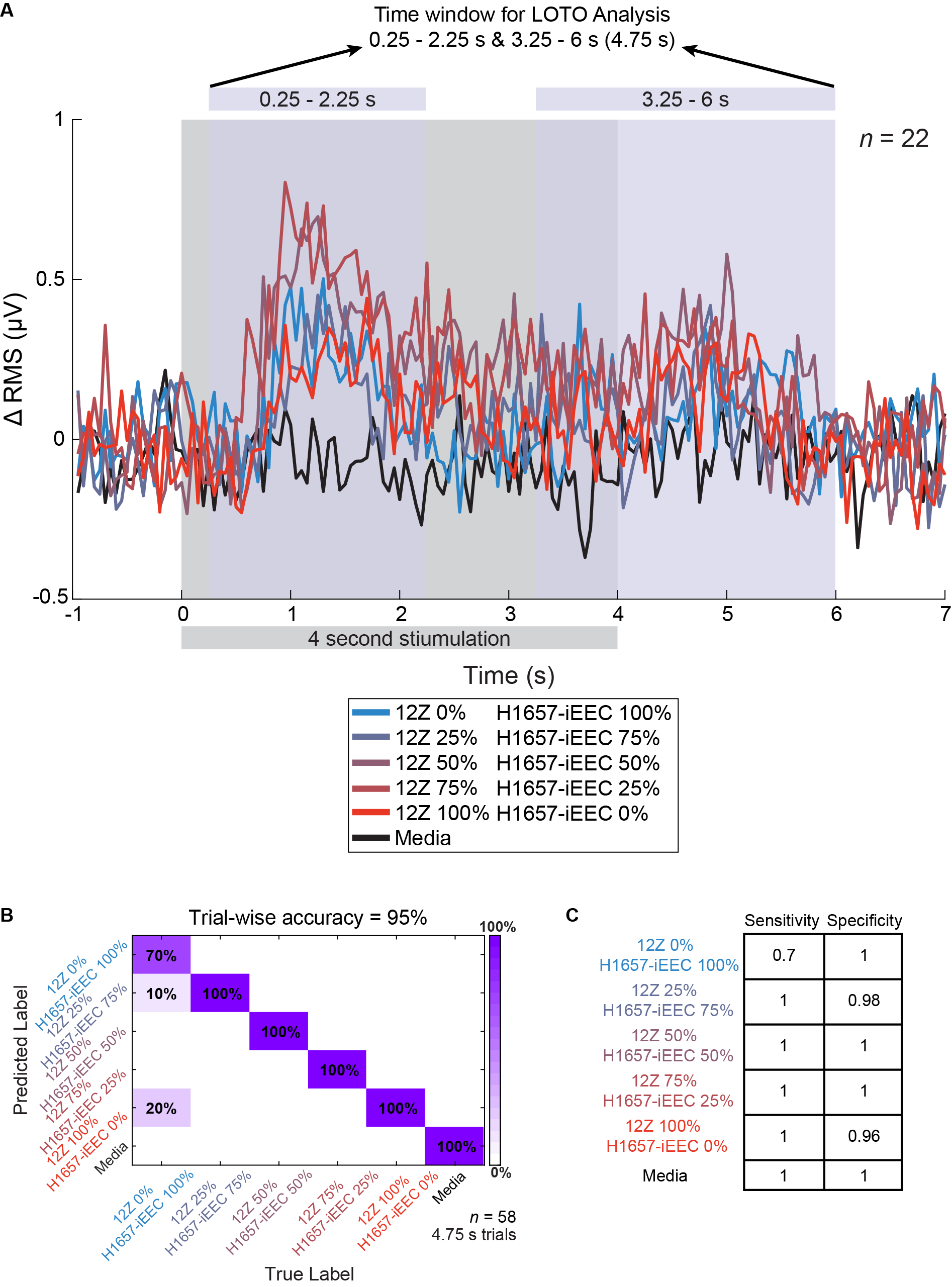

**Supplementary Figure 11. Classification of co-cultures via the neural “on” and “off” responses from emitted gas mixtures.** (**A**) Population based PSTH is shown for all 22 electrophysiological recordings after RMS processing for each cell line co-culture. Time labels along the X-axis are relative to the stimulus onset time. The grey box indicates the 4 second stimulus presentation window. The blue boxes indicate the time window durations (encompassing the “on” and “off” response) used for LOTO analysis shown in Fig. 5D. (**B**) Confusion matrix summarizing the high dimensional LOTO analysis for the classification of 4.75 second trials for the ‘on’ response time window of 0.25 – 2.25 seconds after stimulus onset in combination with the ‘off’ response time window of 3.25 – 6 seconds after stimulus onset (see (A)). (**C**) Sensitivity and specificity table summarizing the results from the confusion matrix in panel C. 58 neurons were spike sorted (see Methods) from 22 electrophysiological recording locations for analysis in (B).

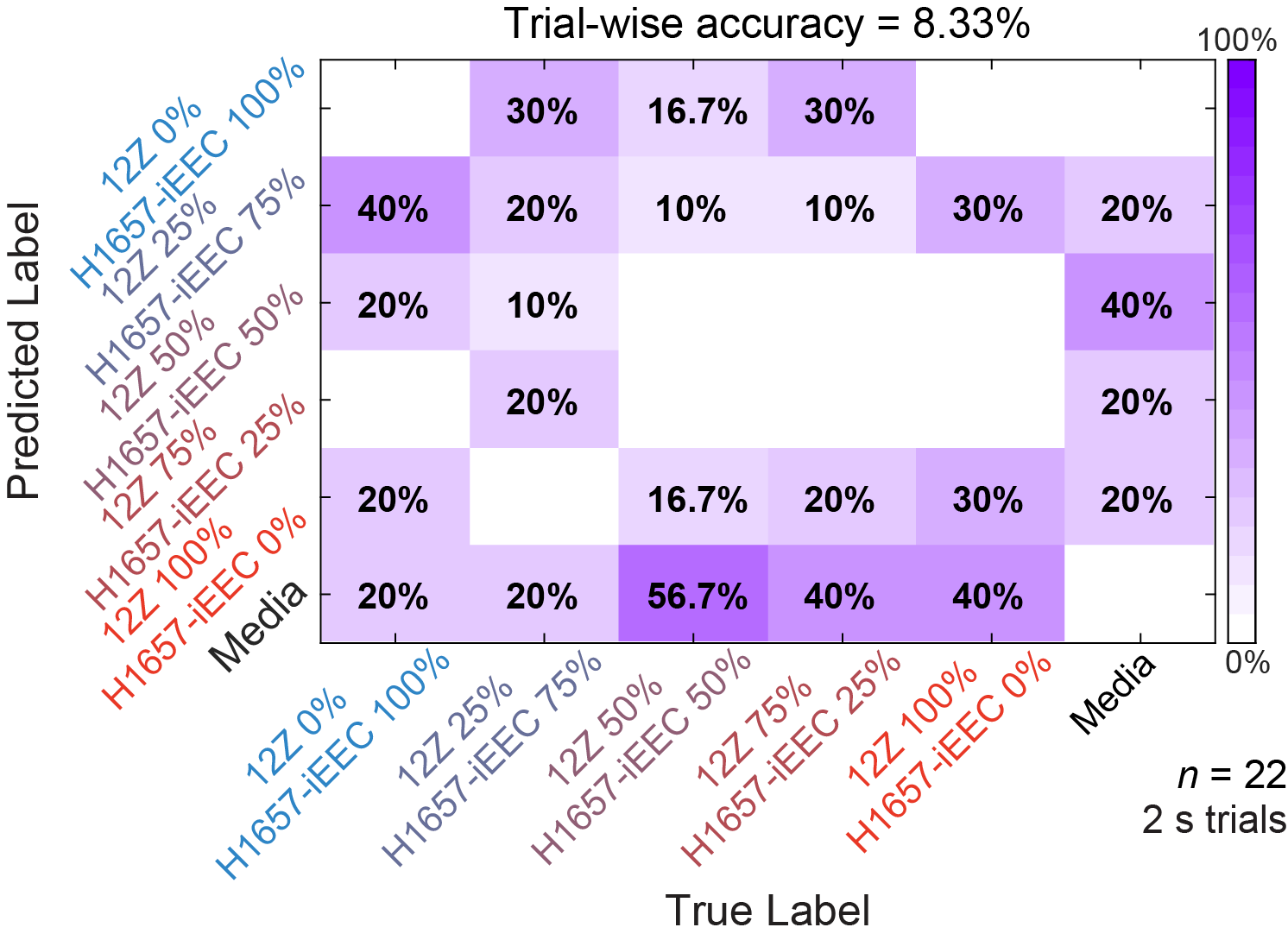

**Supplementary Figure 12. Confusion matrix summarizing the LOTO analysis for the two seconds prior to stimulus onset for cell co-culture.** Confusion matrix summarizing the high dimensional LOTO analysis for the classification of 2 second trials for before the stimulus onset. Data was RMS filtered (see Methods) from 22 electrophysiological recording locations for analysis.

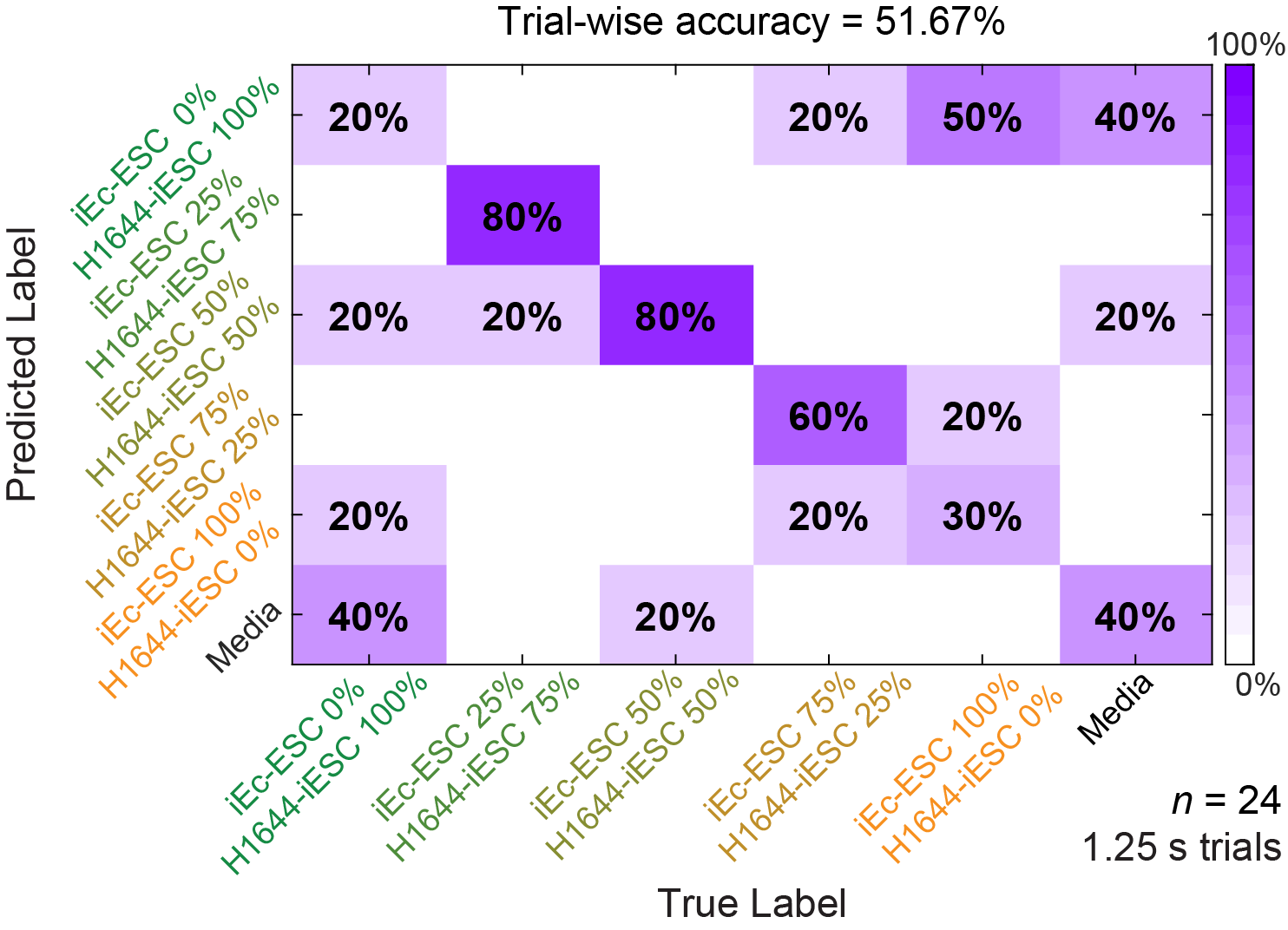

**Supplementary Figure 13. Stromal individual cell line and co-culture classification analysis.** Confusion matrix summarizing the high dimensional LOTO analysis for the classification of 1.25 second trials for the time window of 0.5 – 1.75 seconds after stimulus onset. Data was RMS filtered (see Methods) from 24 electrophysiological recording locations for analysis.

**Supplementary Table 1. Comparison of neural data processing techniques.**

| **Spike Sorting** | **RMS Filtering** |
| --- | --- |
| - Manual process requiring supervision from a skilled researcher - Time intensive - Many methods/algorithms exist^84^ - Variability of human spike sorting performance has been previously noted^75,76^ - Single-neuron activity is obtained - Established analysis for estimating neural population dynamics - Eliminates signals that do not pass statistical tests for inclusion into the dataset | - Automatic and unsupervised processing - Rapid and fast filtering of neural responses - Computationally inexpensive - No variability of processing performance - Single unit activity cannot be obtained - Established mathematical equation for signal processing^85^ - Represents the average power of a waveform - Uses the total energy of the signal obtained from each electrode for inclusion into the dataset |
